# One cell, one drop, one click: hybrid microfluidic mammalian single-cell isolation

**DOI:** 10.1101/2020.01.24.908202

**Authors:** Kenza Samlali, Fatemeh Ahmadi, Angela B.V. Quach, Guy Soffer, Steve C.C. Shih

## Abstract

The process of generating a stable knockout cell line is a complex process that can take several months to complete. In this work, we introduce a microfluidic method that is capable of isolating single cells, selecting successful edited clones, and expansion of these isoclones. Using a hybrid microfluidics method, droplets in channels can be individually addressed using a co-planar electrode system. In our hybrid microfluidic device, we show that we can trap single cells and subsequently encapsulate them on demand into pL-sized droplets. Furthermore, individual cells inside the droplet can be released from the traps or merged with other droplets by simply applying an electric potential to the electrodes that is actuated through a user interface. We use this high precision control to sort and to recover single isoclones to establish monoclonal cell lines, which is demonstrated with a heterozygous NCI-H1299 lung squamous cell population resulting from loss-of-function eGFP and RAF1 gene knock-out transfections.

## 1. Introduction

Gene editing in mammalian cells has become accessible and increasingly less time consuming due to the availability of new editing tools that allow for rapid and precise edits. Using improved versions of CRISPR-Cas9^[1,2]^ along with better methods to control the cell’s double-strand break (DSB) repair mechanisms,^[3]^ Cas9 has become the driving force to engineer new cell lines. Currently, using Cas9 for gene-editing shows widespread benefits for generating new cellular therapies^[4,5]^ and creating new genetic models for cancer^[6,7]^. To create new edited cell lines, evaluating the properties of single clones (i.e. a single edited cell) is especially important because of biallelic editing differences and non-homologous end-joining DSB repair mechanisms generating indels that differ between individual clones.^[8–10]^ Isolating single clones provides a method for enriching correctly edited cells in which one can correlate the phenotypic changes to a specific clonal genotype that will facilitate downstream characterization.

The process of genome editing mammalian cells typically consists of *in silico* design of the guide, cloning the guide into an expression vector, transfection, selection, sorting, and expansion of homogeneous clonal lines.^[11]^ Currently, the design of the guide and the act of transfecting cells can be done in less than a day.^[11]^ With the advent of new automation tools and methods continuously being developed, the process of synthesis^[12]^, assembly^[13,14]^, and transfection^[15–17]^ are becoming, faster, cheaper and more efficient. However, selection and enrichment of transfected clones (especially from knock-out experiments), and the sub-cloning of sensitive cell lines (e.g, hPSCs) still faces low efficiencies. Current methods to isolate single clones are to use limited dilution or colony picking to separate single isoclones and to generate a homozygous progeny.^[18,19]^ The laborious and time-consuming process, the high dilution requirements, and inherent probabilistic nature for limited dilution are not ideal for increasing the chances for obtaining a single clone. Fluorescence-activated cell sorting (FACS) and colony pickers can provide a method to generate clonal cell populations but are associated with high infrastructure and maintenance costs, requires frequent downstream optimization, and usually require a large starting cell population. Furthermore, these physical methods have also been shown to be damaging and induce stress-response in cells.^[20,21]^

Droplet-based microfluidic systems are ideal systems for single-cell manipulation and analysis. These biocompatible systems mimic the physics of the cellular environment and in doing so, they reduce the physical stresses often exerted on cells by traditional tools or automation such as FACS or colony pickers. They are also typically low in infrastructure and operational costs and operate under much lower volumes (∼pL range).^[22–25]^ Some groups have already addressed multiple steps in the gene editing pipeline using microfluidics, including mammalian cell culture ^[26,27]^, transfection of mammalian cells^[28–36]^, as well as the sorting or selection of transfected mammalian cells^[37]^. Droplet-in-channel microfluidics can operate in ultra-high throughput ranges while generating single-cell containing droplets.^[38–41]^ A pitfall with these systems is that it is difficult to manipulate and to control the droplets in parallel. Digital microfluidic systems (DMF) can alleviate the challenges associated with droplet-in-channel systems since DMF are able to address each droplet individually and having this control is especially useful in multi-step procedures such as transformation and enzymatic assays^[42,43]^, drug and inhibitor screening^[44]^, and gene-editing^[27]^. Recently, we have combined both of these platforms together, which we call a ‘hybrid’ microfluidic platform, placing co-planar electrodes (i.e. ground and activated electrodes on the same plate) under microfluidic channels to have individual control of the droplets in channels.^[45]^ Given the increased control of the droplets that hybrid microfluidic technologies have shown^[46–51]^, there is an opportunity to use this technology as a method to control the isolation of mammalian isoclones.

Here, we developed a deterministic ‘one-droplet-one-cell’ hybrid microfluidic system that can trap single isoclones and subsequently and deterministically place them in individual droplets. These single cell containing droplets can be released from traps in two directions, kept in position, or have the opportunity to be merged with other droplets, allowing this device to be used for various manipulations of the individual clones. To show the versatility of our device, we have shown its ability to establish isoclonal mammalian loss-of-function cell lines from gene knock-out experiments by sorting and recovering transfected clones of a NCI-H1299 lung squamous cell carcinoma.

## 2. Results and Discussion

### 2.1 The design of a hybrid microfluidics system for single-cell manipulations

Figure 1 depicts the representative device used for single-cell trapping and for gene-editing assays. As shown in Figure 1A, a microfluidic device consists of three layers: a patterned electrode layer, a 7 µm SU-8 5 dielectric layer and a PDMS-based channel layer consisting of 35 μm height and a main channel width of 50 μm. The ‘hybrid’ microfluidic device consists of a bottom digital microfluidic layer (i.e. electrode and dielectric) along with a top patterned channel layer in which droplets are manipulated in an oil phase.^[45]^ The device is divided into two sections: 1) an on-demand droplet generator and 2) a single-cell droplet array. As shown in Figure 1B, the on-demand droplet generation consists of co-planar electrodes that will actuate the aqueous flow (using electric potentials) to the orthogonal continuous oil flow that will break the continuous aqueous flow into discretized droplets under the control of the user. The single-cell droplet array (Figure 1C) can trap single cells, after which they are encapsulated in droplets that are generated within the traps by the application of an electric field. The device contains 12 traps (six are equipped with electrodes) in which a single cell is displaced into an empty trap. Tubing connects the two parts of the device to transfer the droplets from the droplet generator to the single analysis part of the device. The device contains two inlets: (I1) oil and droplets and (I2) cells and priming and two outlets: (O1) waste and (O2) sample recovery and flow reversal (Supplementary Figure 1).

**Figure 1.**
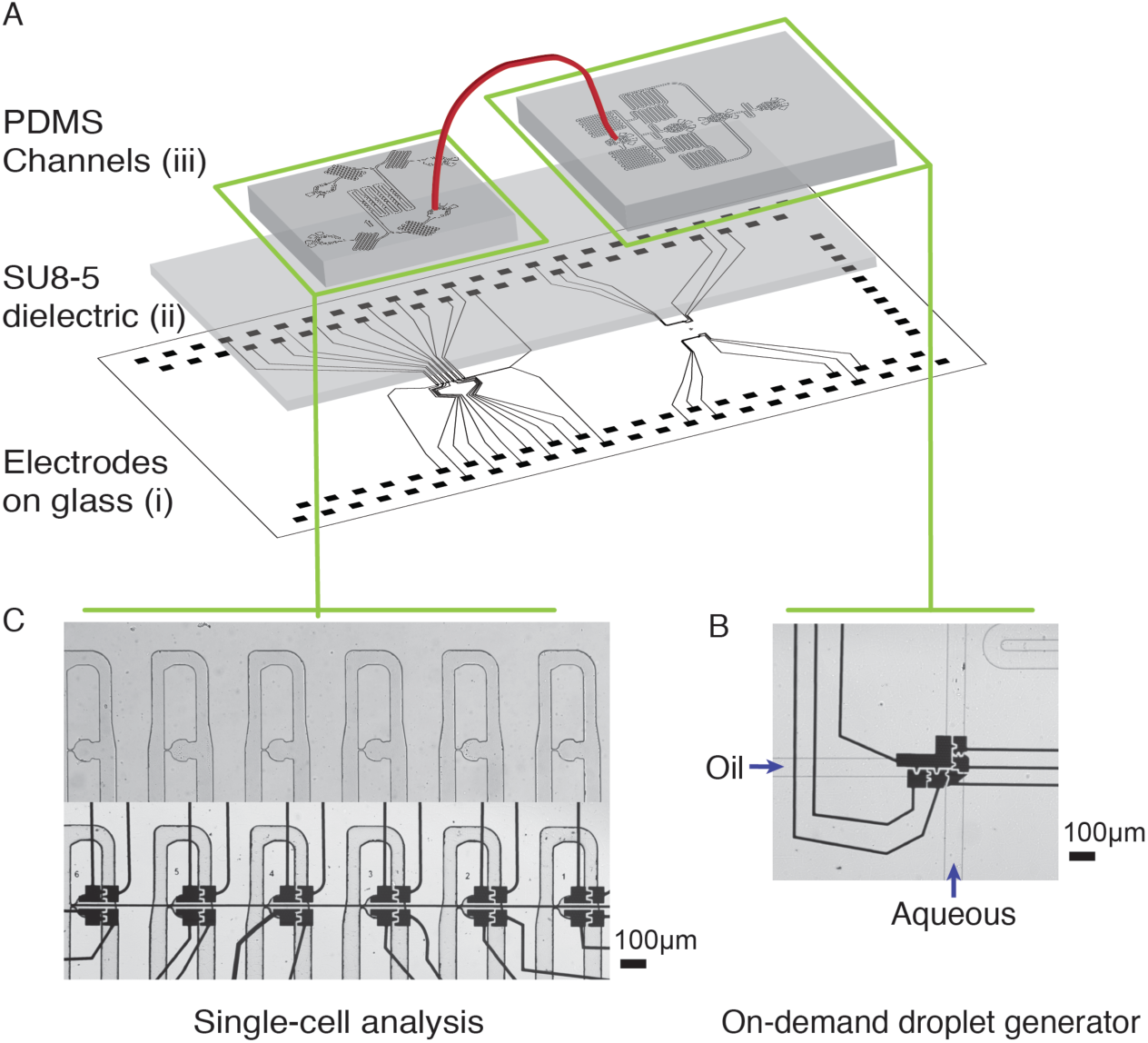
Integrated droplet digital device for on-demand single-cell encapsulation and single cell analysis. The three-layer device consists of a digital microfluidic layer with chromium electrodes patterned on glass, a 7 µm thick SU-8 5 dielectric layer and a PDMS channel layer of channels of height 35 µm and width of 50-75 µm. B) The droplet generation device contains two T-junction droplet generators, under which several electrodes are located below the channel to initiate on-demand droplet generation. C) The single-cell analysis device contains two inlets, and two outlets. The trapping area contains cell traps with 8 µm constrictions and four electrodes below the trapping area. More details related to the device and channel, electrode and wiring sizes is shown in Supplementary Figure 1.

In designing the system shown in Figure 1, there is a design element that requires consideration for reliable cell trapping and encapsulation. Two commonly used methods for single cell isolation – trapping and encapsulation – are complementary. The traps are designed such that they could trap single cells with high efficiencies, yet also allow for a smooth phase change. We followed resistance based design guidelines as reported for microfluidic serpentine trap designs for droplets ^[52–54]^ and cells ^[55–57]^. The design element concerns the location of the traps relative to the main channel such that both single cells trapping, and phase change can occur. By modeling flowrate profile and velocity streamlines, we optimized the channel geometry (Supplementary Figure 2), such that volumetric flow rate through the trap (Q_trap_) is greater than volumetric flow rate through the bypass channel (Q_bypass_) when there is no cell in the trap. We found that positioning the trap near the curvature of the main channel (i.e. the end of a serpentine channel) along with a narrow (∼50 μm) trap entrance and a narrow (∼50 μm) width of the main channel immediately after the trap opening, prevented cells bypassing the empty traps. The optimized placement offers a higher effective hydrodynamic resistance in the bypass channel (R_bypass_) than through the trap (R_trap_). Hence, the flowrate in the trap is higher compared to the bypass channel (Q_trap_ > Q_bypass_) to maintain the same pressure drop (as shown from other studies ^[58]^). Furthermore, the design offers two additional advantages: (1) if a cell is trapped, it is unlikely for another cell to flow into the same trap since this increases the R_trap_ (and reduces Q_trap_) and (2) during a phase change for single-cell encapsulation (trap based droplet generation) (described below), the resistance in the trap is sufficiently higher than the bypass channel which will help preventing squeezing the cells out of their traps. A mathematical description and the simulation details are described in Supplementary Note 1 and 2.

Finally, we also note that one of the main goals of this work is to automate the process of actuation and droplet manipulation, but a key challenge is to integrate and to control the multiple pieces of hardware into one software framework. In this work, the microfluidic device is connected to two main hardware components: the in-house automation system (i.e. optical switches) and a syringe pump system (Supplementary Figure 3, Note 3). The automation system serves the purpose to provide electrode actuation and impedance feedback and the syringe pump system is to control the flow rates in the device.^[43,59]^ Since these two hardware systems are operating on different software protocols, we developed our own Python based framework with a simple user interface for one-click droplet operations.

### 2.2 Single cell trapping and droplet encapsulation

Figure 2 illustrates the optimized device operation procedures for trapping single cells and encapsulating the cells inside droplets using a ‘hybrid’-based microfluidics platform. As detailed in the Experimental Section, the device operation procedure consisted of priming, cell loading, phase change, and encapsulation followed by droplet release (see Supplementary Note 4 regarding details on device assembly and channel treatment prior to operation). The highlights of this procedure are represented as a schematic, shown in Supplementary Information Figure 4. First, devices were primed with 2% Pluronics F-127 for at least 5 min to decrease cell adhesion to PDMS. Second, an aqueous flow containing fresh media with mammalian cells (MCF-7 breast cancer cell line) was introduced into the trapping device at a concentration of 10^5^ -10^6^ cells mL^-1^ (see Figure 2A for an image of six individually trapped cells). We evaluated the efficiency of cell trapping as a function of flow rate. Using our design, the optimal range of flow rates to trap individual cells is between 1 – 4 nL s^-1^ (Figure 2B). At this range of flow rates, cells are unlikely to occupy traps with multiple cells and the MCF-7 cells do not squeeze through the traps. Single cells are most efficiently trapped (∼88.3 %) at 5 nL s^-1^ -an efficiency similar to previous studies which required multi-layer displacement structures ^[60]^ or other external forces ^[61,62]^ to trap cells. We also verified single cell trapping by observing 54 consecutive cells passing an occupied trap and all of the cells passed the occupied trap (i.e. the cells remained in the main bypass channel). The conditions defining the success at these flowrates are (1) the optimized channel flow velocity profile and slanted overhang along the main channel (close to the trapping region) to steer the flow towards the trap (2) designing a 8 µm constriction which is smaller than the cell size, and (3) the physical properties of MCF-7 cells (i.e. lower deformability).^[55,58]^

**Figure 2.**
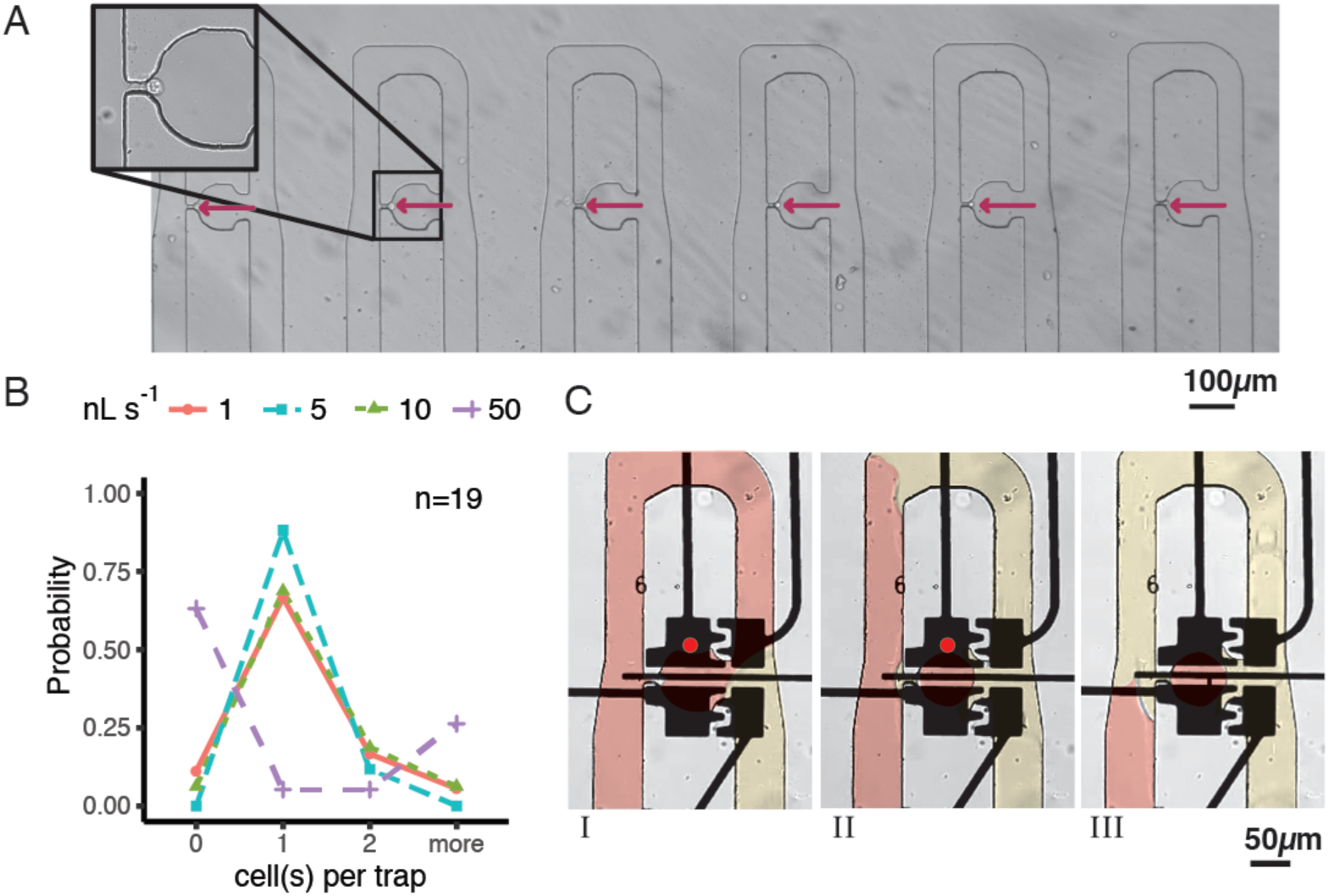
Cell trapping and encapsulation. A) Single MCF-7 cells trapped in the constriction. An aqueous PBS flow is present (a bright field, 4X image). B) Efficiency of trapping cells at different flowrates (1, 5, 10, 50 nL s^-1^). The cell concentration was kept constant at 5 x 10^5^ cells/mL in PBS and the likelihood of capture (probability) was measured at 10 min. C) Images showing the encapsulation procedure. Frame I: A single MCF-7 cell trapped. A HFE 7500 and 2% RAN surfactant was loaded into the device at 4 nL s^-1^. The trap electrode was actuated (15 kHz, 126 V_RMS_). Frame II: A droplet was formed within the trap and the oil phase continues through the bypass channel. Frame III: An encapsulated MCF-7 cell in a droplet. False colour was used to highlight the liquid flow and droplet inside the channel.

Following trapping of the cells is the encapsulation of a single cell inside a droplet using a phase change procedure. Popular passive single-cell encapsulation is known to be a procedure that follows Poisson statistics, generating droplets with none, one or more cells.^[63]^ Using the hybrid device, we can generate a droplet within a trap on-demand, and thus encapsulate the trapped cells deterministically. The trapping and encapsulation are done by (1) flowing an oil phase through the entire channel and (2) applying an electric potential to the electrodes below the trap when the oil flow approaches the trap. Figure 2C shows three images taken from video frames showing the encapsulation process. Four coplanar electrodes (size = ∼100 μm, area = 0.06 mm^2^) were used for the encapsulation event – two electrodes below the main channel and two electrodes below the trap. In Frame I, all electrodes are grounded. An oil flow enters the main channel for the purpose of a phase change. In Frame II, the electrode below the trap is activated while the other electrodes are grounded. The aqueous phase and the single cell remain inside the trap when the oil flow (in the main channel) “cuts” the aqueous phase at both ends of the trap. Generated cell containing droplets are on average 150.3 ± 5.6 pL in volume (N = 11) (see Supplemental Information Table S1 for additional device statistics). In Frame III, all potentials are grounded, and the oil phase flow continues to the next trap to perform the next encapsulation procedure. To aid the design of the trap and to determine the optimal actuation sequence, we have simulated the electric potential and electric field distributions (Supplementary Figure 5, Figure 6). As shown, the electric field density (∼ 5 x 10^6^ V m^-1^) is induced between the main channel and the trapping region. The field gradient induces an electrostatic force that will pull the liquid towards the trap (similar to digital microfluidic droplet actuation ^[64]^). Given this capability, the device has means to encapsulate cells in droplets on-demand without Poisson-based statistics. The details of the simulation are described in the Supplementary Note 2, Table S2.

Given its feasibility, the success of trapping and encapsulation is highly dependent on device fabrication and operation methods. For example, the reliability of electrode actuations and resulting droplet operations heavily depends on the alignment of the electrodes and channels. To minimize the difficult task of alignment, we used the ground wire and the gap between electrodes to serve as an alignment mark. Furthermore, we divided the device into two components (droplet generator and serpentine trapping channel) to fit the features within the view field of the microscope and to minimize PDMS shrinkage.^[65]^ Next, we and others, also observed that alignment has to be performed quickly^[66]^ due to the temporary effects of oxygen plasma on PDMS and glass. Additionally, the process of inserting and removing tubing from the inlets and outlets requires slow insertion and removal since quick removal usually causes air bubbles inside the channels. Air bubbles should be avoided while changing from priming solution to cell solution and when priming the channel with the oil flow since the solutions can block flow inside traps and can push cells out of their traps or disturb the stability flow causing irregular flowrates inside the channel. The air bubbles also generate pressure differences inside the channel which may lead to droplet breakup and movement. Our solution is to insert and to remove the tubing gently from the inlet and outlet respectively while maintaining high flow rates to minimize large pressure differences in the main channel. Finally, it is important to perform a thorough cleaning of the traps by removing the remaining oil emulsions in the 8 µm trap constrictions to ensure high cell trapping efficiency for the next set of trapping experiments.

### 2.3 On-demand droplet generation, release, and keep

After trapping and phase change, we turned our attention to other droplet operations such as droplet generation or keeping and releasing the droplets containing single cells. Generally, in droplet-based microfluidic devices, controlling droplet positions inside the channels is performed by using passive structures ^[54,55,58]^, valves^[67,68]^, or external forces (optical, acoustic, dielectrophoresis) ^[54,69,70]^. For example, Sauzade et al. uses serpentine channels containing droplet traps under forward flow to trap droplets and uses reverse flow to hydrodynamically release droplets.^[55]^ The platform presented here can perform multiple droplet operations, such as a trapping operation under forward flow, release operation under forward/reverse flow, and keep operation under reverse flow. Our device have no additional channel structures that have been fabricated and there is no reliance on timing to control the droplets (as required by other works ^[71,72]^). The main contributor to controlling the droplets on our device is the application of electric potentials to the electrodes (similar to digital microfluidic systems ^[73]^) such that the above-mentioned operations can be performed with high fidelity.

To characterize releasing operations, we have tested the likelihood for droplet release at different flow rates (for the forward and reverse flow directions) using electric potential or via pressure-driven flow. Figure 3A (Frames I-IV) shows the actuation sequence for releasing a droplet under forward flow. The droplet is released by actuating electrodes below the trap (Frame II) followed by activating an electrode below the main channel and the trap (Frame III). By using this specific sequence, the electric field density directs the droplet from the trap towards the main channel in the direction of the flow (Supplementary Figure 6). We also tested the likelihood for droplet release at different flow rates in forward direction (from inlets to outlets) (Figure 3B). As shown, low forward flow rates (< 1 nL s^-1^) give rise to high probability (> 95 %) of being able to release the droplet. Since droplets are trapped due to the hydrodynamic pressure, *P_h_*, of the oil flow and the droplet is controlled by using electrostatic forces (*F_elec_)*, droplets can be released when the electrostatic force *F_elec_* is greater than the *P_h_* generated by the flow in the main channel. This relationship also holds true when there is no flow rate applied. In this case, the droplet is released from the trap but is static at the entrance of the trap since there is no flow. While without any electrostatic force (i.e. no electric field applied) at any given flow rate, the droplet is never released from the trap.

**Figure 3.**
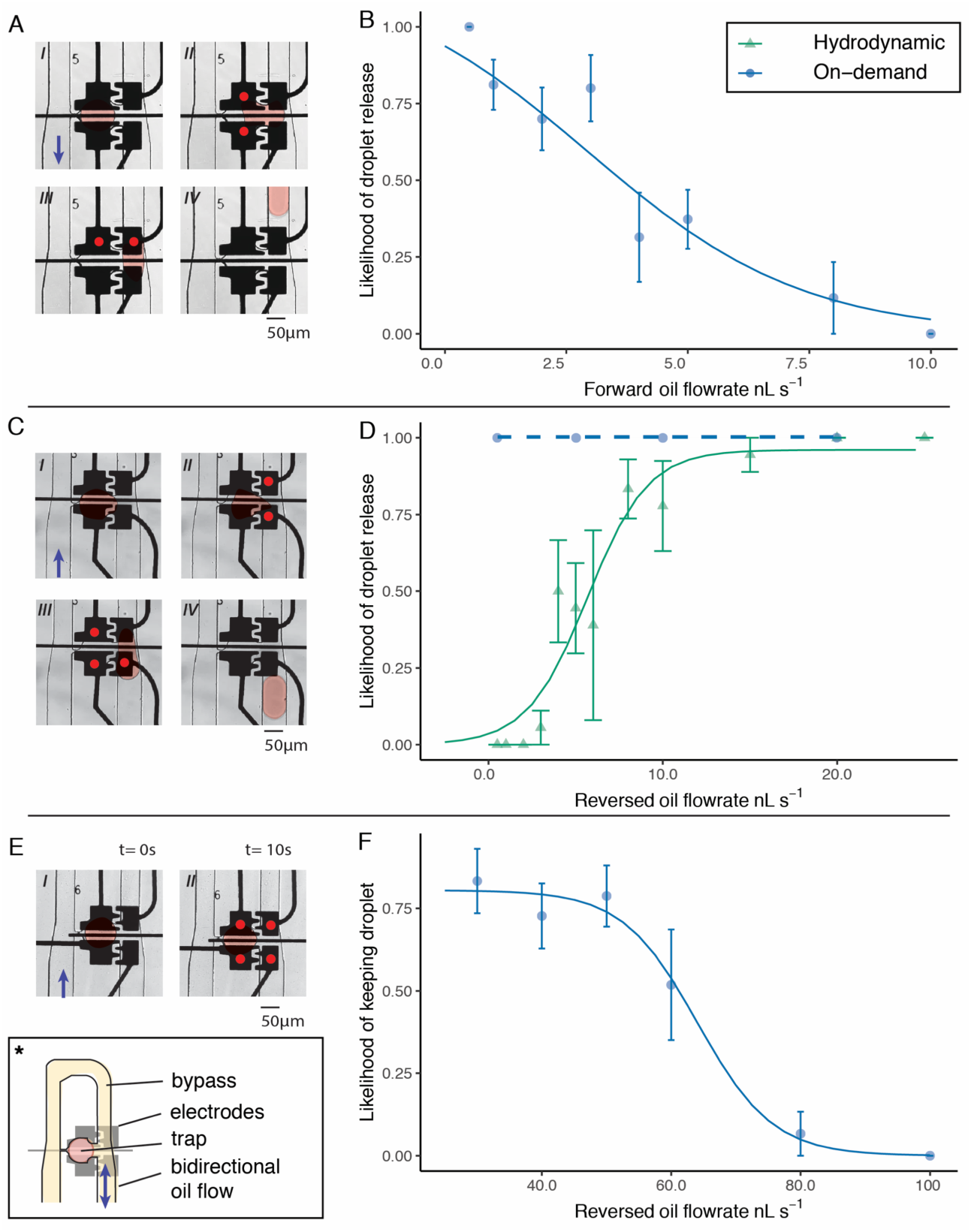
On-demand droplet operations. Actuation patterns are indicated with a red dot (bright field, 15X). Image showing the bypass channel, trap and flow is indicated by a (*). False colour was used to highlight the droplet. A) Actuation sequence of releasing droplet towards outlet on demand (15 kHz, 126 V_RMS_). B) Efficiency of release of droplets under forward flowrate, under increasing flowrates (n = 8, 10 replicates per trap). C) Actuation sequence of releasing droplet towards inlets on demand (15 kHz, 126 V_RMS_). D) Efficiency of release of droplets under reversed flowrate. Hydrodynamically, droplets are released more efficiently towards the inlets, under increasing flow rate. With on-demand release, droplet show near-perfect release with applied flow rates between 0.5 - 20 nL s^-1^. E) Actuation sequence of keeping droplets within trap under reversed flow rate (15 kHz, 126 V_RMS_, 10 s). F) Efficiency of keeping droplets on-demand under reversed flow rate. Droplets can be kept efficiently (∼78 %) for flow rate lower than 45.4 nL s^-1^.

**Figure 4.**
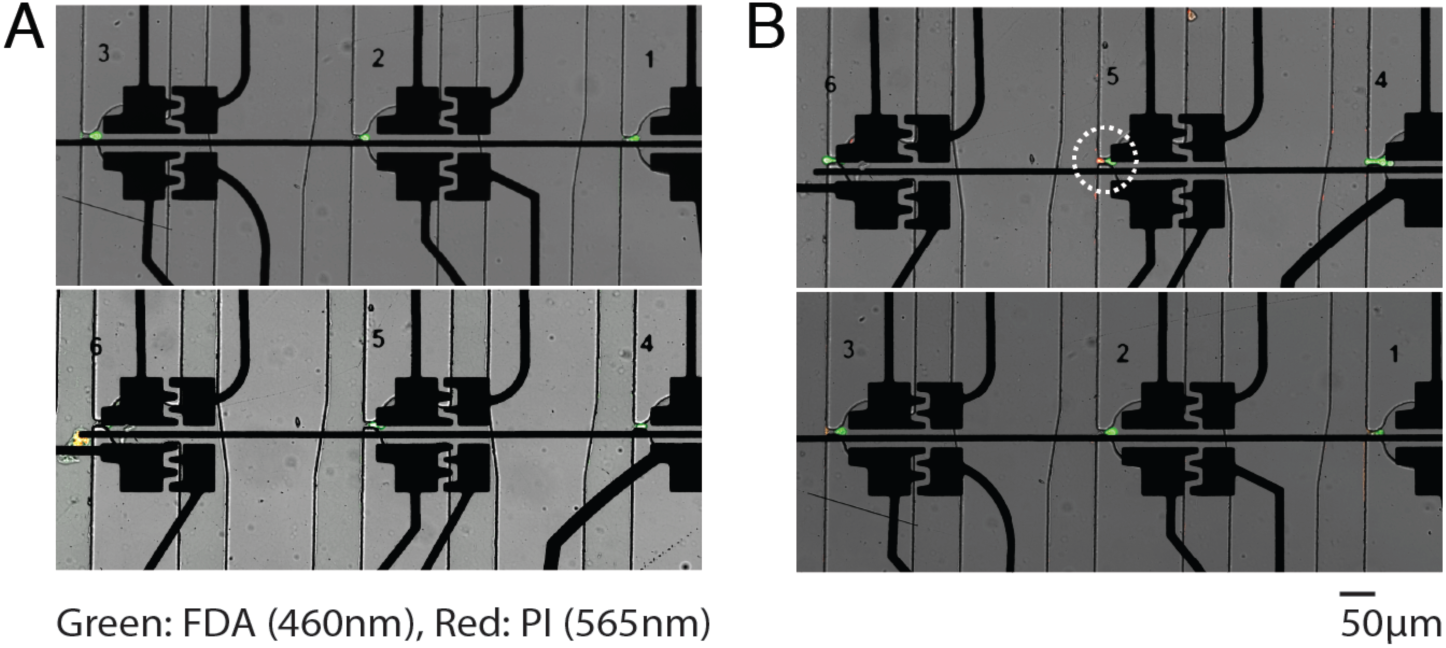
Viability assay of MCF-7 cells. A) Trapped MCF-7 cells. FDA stain reveals live cells and PI stain reveals dead cells. B) Trapped MCF-7 cells stained with FDA/PI after actuation of electrodes for 20 s (15 kHz, 126 V_RMS_) and 10 min incubation on device. No significant difference in viability can be detected after actuation of electrodes.

Next, we tested the likelihood of releasing droplets with reverse oil flow, with and without on-demand actuation. As shown in Figure 3C (Frames I-IV), the actuation sequence under reverse flow (from outlets to inlets) is similar to the actuation sequence for the droplet release under forward flow. In contrast to with forward flow, the probability of releasing a droplet is most likely to occur at higher flow rates (> ∼10.9 nL s^-1^) (without actuation; hydrodynamic flow only). The lower flow rates are more likely to keep the droplet inside the trap (Figure 3D) – a similar trend observed in other studies.^[74]^ When actuation is implemented, the droplet can be released from the trap at any time and there is no dependence on the reverse flow rate using a specific actuation pattern. This is an exciting result since this enables the user to release and to select droplets on-demand and in parallel without the need for dielectrophoretic, acoustic or magnetic sorting techniques. Thus, this represents a significant advance over other droplet-based microfluidic systems that implement trapping and releasing droplets.

In some cases, keeping droplets inside a trap is also a desired operation (as shown from these studies^[45,75,76]^). Figure 3E are images showing the actuation sequence for keeping a droplet. Four electrodes are activated to ensure the highest electric field density is centered at the opening of the trap to prevent the droplet from escaping into the main channel (Supplementary Figure 6). The likelihood of the droplet being kept in the trap (∼77.6 %) is when flow rates below 45.4 nL s^-1^ are applied in the main bypass channel (> 95% logistic regression model asymptote) (N = 10) (Figure 3D). The main reason for this trend is that above 45.4 nL s^-1^, the hydrodyanamic pressure in the channel is greater than the electrostatic force from the applied potential (*P_h_ > F_elec_)*. It is possible to increase the applied potential (> 126 V_RMS_) to the electrodes (to increase the electrostatic force and work under higher flow rates), but this may induce dielectric breakdown^[77–79]^, followed by electrolysis or Joule heating which can ultimately lead to cell stress and to changes in genomic regulation in cells^[80]^. Hence, for gene-editing experiments discussed below, we used flow rates below 45 nL s^-1^ to keep the droplets inside the trap while maintaining applied potentials below 126 V_RMS_.

Similar to our previous work^[45]^, we generated droplets on-demand to add reagents to other droplets inside the device. We performed on-demand T-junction droplet generation (Supplementary Figure 8). At oil flow rates lower than 2 nL s^-1^ and higher than 2.5 nL s^-1^, on-demand droplet generation became difficult, due to the inability of the oil flow to shear off a droplet at low flowrates, or the pressure of the oil flow being larger than the electrostatic force. Droplets arriving through the droplet bridge in the trapping device, remain the same volume as droplets made in the droplet generator device (unpaired two-sample T-test, P > 0.05). As shown in Supplementary Information Figure 9, we can merge incoming droplets with trapped droplets on demand. After forward releasing of a droplet, we were also able to merge two droplets from different traps, each containing a single cell. An advantage of on-demand merging^[81]^ is that it does not rely on the tedious synchronization of two streams of droplets for droplet coalescence or does not require any pressurized channel.^[46,72,82–86]^ In addition, the high variability in droplet volumes may interfere with biological assays, however, both generating droplets on-demand with a T-junction as generating single-cell containing droplets by phase change, showed low variability in the droplet volume (250.9 ± 39 pL and 150.3 ± 55.6 pL respectively) (Supplementary Figure 8). To our knowledge, the compendium of results in Figure 3 represents the first report that describes multiple on-demand droplet manipulations with individual and parallel droplet control in droplet-based microfluidic devices. These droplet operations in addition to the deterministic encapsulation provide a powerful device for sorting or assays on individual isoclones.

### 2.4 MCF7 cells are viable after electrode actuations on hybrid device

As previously described, our device can perform a variety of droplet operations. We tested our system and all of the above operations with MCF-7 human breast cancer cells to evaluate the capability to precisely control trapping and encapsulation of single mammalian cells. Following the procedure of trapping single-cells in our system, single MCF-7 cells have been trapped and their viability have been measured while being trapped. As shown in Figure 4A, we performed a viability assay by flowing a solution of fluorescein diacetate (*λ*_ex_: 490 nm, *λ*_em_: 526 nm) and propidium iodide (*λ*_ex_: 488 nm, *λ*_em_: 617 nm) through the channel to label live and dead cells respectively (which happens after priming the device and trapping of single cells). In our experiments, we observe a small loss of viability and hypothesize that this is attributed to cells being outside their *in vitro* culturing environment – e.g., the resuspension of the cells in PBS and injecting the cells into a syringe. We also tested the cells under a different condition (applying a potential for 20 s at 15 kHz and 126 V_RMS_) and observed a similar viability (∼ 87.5 %) (Figure 4B). This is an expected result since it is previously shown that the actuation of electrodes in digital microfluidics and the shear stresses in channel microfluidics do not affect the viability of commonly used mammalian cell lines.^[26,80]^ Yet, increasing potentials ^[80]^ or squeezing cells ^[31]^ may upregulate different types of stress genes^[80]^, but in this study we avoided very high potentials (> 140 V_RMS_) and squeezing the cells.

### 2.5 Successful recovery and expansion of single-cell isoclones from a heterozygous engineered cell population by single-cell manipulation on device

Previously, other groups have recovered droplets from a microfluidic chip on a substrate to recover cells using a chemical emulsion breaking method^[87]^, centrifugal methods^[88,89]^, or automated dispensing methods^[90,91]^. Typically, these methods are performed on an emulsion of multiple droplets which makes it difficult to recover a single drop or cell and also lowers the viability of the recovered cell. The deterministic encapsulation and on-demand release of droplets in our platform enabled a high recovery rate (near perfect) that is capable of retrieving the content of a single droplet from a water-in-oil microfluidic emulsion into a single well of a 96-well plate (see Supplementary Figure 10 for a detailed workflow). A dual-labeled NCI-H1299 lung squamous cell line expressing eGFP have been used as a model to facilitate the optimization of a single cell recovery from the hybrid device. Following an on-demand forward release of a single isoclone, we transferred the droplet to a hydrophobic PTFE membrane to absorb the HFE 7500 oil while exposing the droplet containing a single cell (a modified protocol from a previous study^[92]^). By rinsing the emulsion with FC-40 oil, the emulsion destabilizes and releases the isoclone into a droplet containing fresh cell culture media which is subsequently transferred to a 96-well plate. We tested this procedure using the NCI-H1299 eGFP expressing cell line and we were able to perform the trapping, encapsulation, isoclone release, and recovery of the isoclone in less than 1 h. To our knowledge, this is the first time that such a platform can perform these operations in this rapid timeframe. We also transferred the eGFP^+^ isoclone to a well plate for expansion and we observed cell growth after 5 and 7 days (Figure 5).

**Figure 5.**
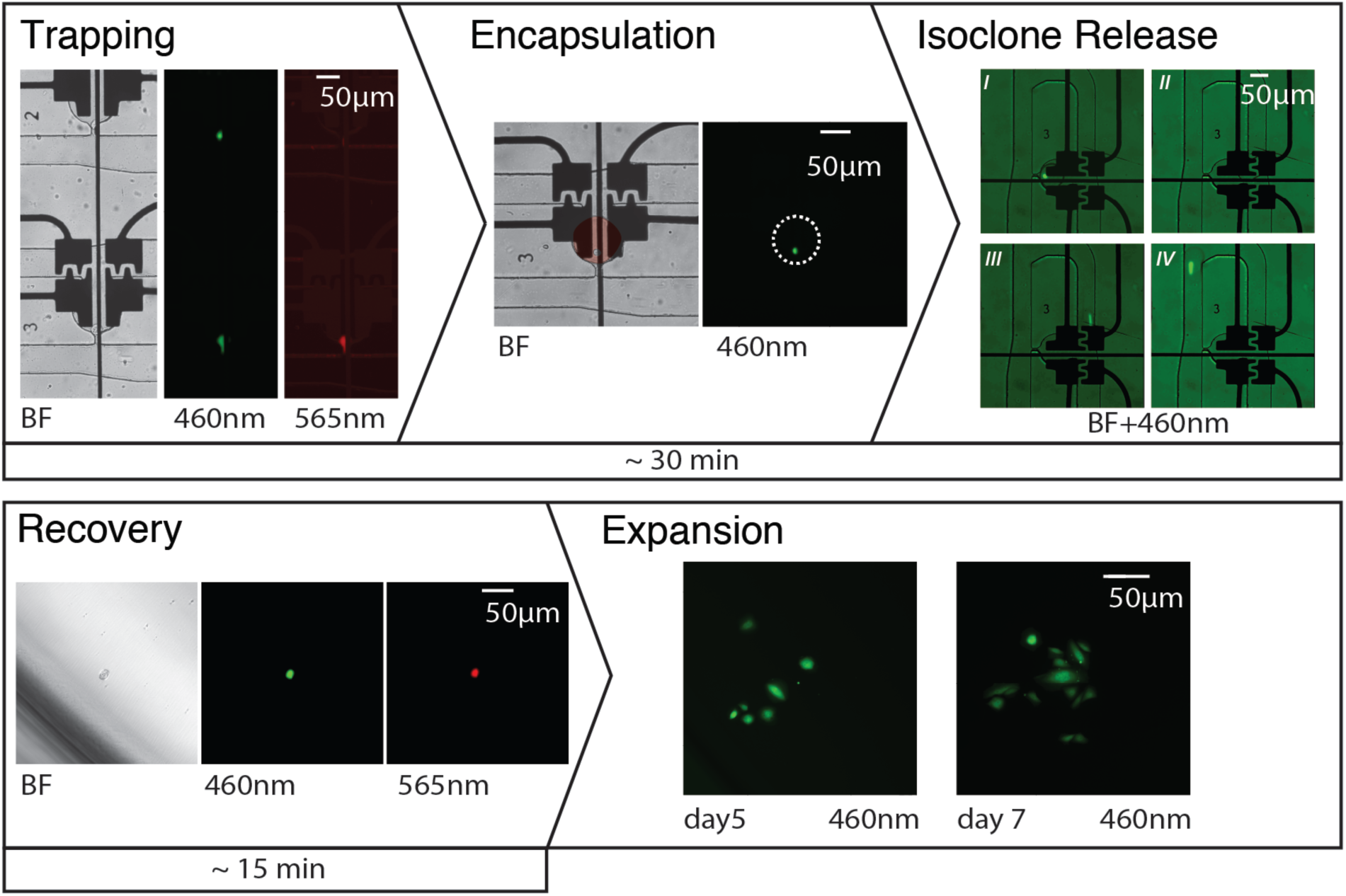
Screening and sorting of edited H1299 isoclone. Bright-field, 460nm and 565nm excitation microscopy images of H1299 cells showing the trapping, encapsulation, isoclone release and recovery, and expansion. Trapping: H1299 cells (eGFP^+^) resulting from lipid mediated transfection were trapped and screened for mCherry expression (red). Encapsulation: A trapped cell show successful encapsulation with a droplet. Isoclone release: The process to release the isclone by on-demand methods (Frames I-IV). Recovery: The droplet containing the isoclonal cell is collected from the outlet into a capillary and recovered into a 96-well plate. Expansion: eGFP+ cells were maintained and grown for at least 7 days. The whole process (excluding expansion) only took ∼45 min.

Next, using a high copy number CRISPR-Cas9 all-in-one plasmid, we performed an NHEJ-based Cas9 mediated gene knock-out. We lipo-transfected the H1299 cell line and knocked out either the eGFP gene or the RAF1 oncogene. From our flow cytometry results, we obtained ∼15.5% and ∼14.8% transfection efficiency, respectively (Supplementary Figure 11). As shown from Supplementary Figure 12, we obtained a heterozygous population which required sorting to obtain the successful clones. We injected the heterozygous population into our hybrid device, after which we imaged the cells and observed whether a transfected clone was trapped. Due to the low transfection efficiencies, we expected only one out of six functional traps to be filled with mCherry^+^ (16.7%). Indeed, there were times when the traps did not fill with a successful knockout clone, however, increasing the flow rate enabled the non-KOed cells to squeeze through the traps such that a new cell can be trapped. After encapsulation of a successful isoclone, on-demand forward release of a single-isoclone was performed to recover the isoclone (Figure 5). Transient expression of mCherry helped to verify the recovery of the transfected isoclone 24 h after the procedure. Currently, we are verifying the monoclonality of the RAF1^-^ and eGFP^-^ populations after expansion by Sanger sequencing of the knockout region.

Overall, the procedure presented here gives scientists the ability to establish monoclonal lines from heterozygous transfected populations, without the need of an alternative selection, sorting, limiting dilution or a clonal picking step. The simple change of an isoclone library and the versatility in choice of screening method (microscopy, fluorescence based) results in a wide sorting spectrum that can be used for low transfection efficiency cell lines.

## 3. Conclusion

The combination of hydrodynamic pressure and electrostatic force offered in the three-layer hybrid microfluidics device was used to control droplets with a single cell. First, we showed that reliable single cell trapping can be followed with a deterministic encapsulation of the trapped single cells. Using a phase change and electrode actuations, the one-phase cell containing aqueous flow can be turned into a droplet based two-phase flow. Next, we have shown excellent droplet controllability and fully characterized the efficiency of droplet generation and bi-directional release and keeping of droplets. All of these operations, including deterministic encapsulation, were performed by a “click” of a button.

We applied this system to sort and to recover gene-edited single cells from a NCI-H1299 non-small cell lung carcinoma cell population. Single cells from the heterozygous knockout population were encapsulated and sorted based on expression of a reporter protein. Next, we developed a methodology for recovery of an isoclone from a single droplet into a standard 96-well plate and demonstrated this for a knockout cell line generation pipeline. Compared to automated systems used for sorting out isoclones in the cell engineering pipeline (e.g., FACS, limited dilution, and clone picking), our system can work with low sample volumes (< 200 µL) and low subpopulation levels (i.e. hard to transfect cell lines). The procedure is fast and gentle on the cells, as by the viability and expansion results. We believe this could be of particular interest for use with hard-to-transfect primary cells or pluripotent stem cells.

In the future, improving alignment, increasing the number of traps, or further automation capabilities of the sorting and recovery system will greatly improve the functionality of the device. We can also envision this multi-functional platform to be used for performing transfection on device, deterministic merging of two populations of single-cell containing droplets, non-binary single-cell sorting, and potentially expansion of isoclonal cultures within droplets. Hence, hybrid microfluidics creates new avenues towards other mammalian cell assays that have previously been difficult to translate to microfluidic platforms.

## 4. Experimental section

### Reagents and materials

Fabrication materials for hybrid microfluidic devices include the transparent photomask (CAD/Art Services Inc., Bandon, OR), S1811 positive photoresist coated glass slides (Telic, Valencia, CA, USA), MF321 developer (Rohm and Haas, Marlborough, MA, USA), CR-4 chromium etchant (OM Group, Cleveland, OH, USA), AZ-300T photoresist stripper (AZ Electronic Materials, Somerville, NJ, USA), <100> Si wafers (Silicon Valley Microelectronics Inc., Santa Clara, CA, USA), and SU8-5, SU8-2035, SU8 developer (Microchem, Westborough, MA, USA). Polydimethylsiloxane (PDMS) was purchased from Krayden Inc. (Westminster, CO) and chlorotrimethylsilane from Sigma-Aldrich. Polylactic acid (PLA) material for 3D printing was purchased from Shop3D (Mississauga, ON, Canada). DI Water had a resistivity of 15 M*Ω* cm^-1^.

Reagents for device preparation include 3M Novec HFE7500 engineering fluid and the surfactant 3M Novec 1720 (M.G. Chemicals, Burlington, ON, CA), 20 g of 5% wt of PEG fluoro-surfactant dissolved in HFE7500 (Ran Biotechnologies, Beverly, MA, USA), Fluorinert FC-40 (Sigma-Aldrich), Pluronics F-127 (EMD Millipore Corp, Billerica, MA, USA), and Triton X-100 (Sigma-Aldrich). All glass syringes were from Hamilton, Reno, NV, USA. All tubing and fittings were acquired from IDEX Health & Science, LLC, Oak Harbor, WA. Glass capillaries were purchased from World Precision Instruments (FL, USA). 0.22µm hydrophobic PTFE membrane was purchased from Thomas Scientific (NJ, USA).

All cell culture and preparation reagents were purchased from Thermo Fisher (Mississauga, ON, Canada). The cell culture reagents include DMEM, RPMI-1640, fetal bovine serum (FBS), penicillin/streptomycin and phosphate buffer saline (PBS) (Ca^2+^/ Mg^2+^ free). The cell viability reagents include Fluorescein diacetate (FDA) (5 µg mL^-1^) and Propidium Iodine (PI) stock solutions (2 µg mL^-1^). For transfection, a Neon Transfection kit (Mississauga, ON, Canada) and Lipofectamine^TM^ 3000 Transfection Reagent, Genomic Cleavage Detection kit were also purchased. EndoFree Plasmid Maxi Kit for plasmid purification was acquired from Qiagen (Toronto, ON, CA).

### Device fabrication and assembly

The photomasks for the hybrid microfluidic devices were designed using AutoCAD 2017, with an electrode design and dielectric layer on a glass slide (50 x 75 mm), and a channel design fitting a 4”-Si wafer. Electrode and dielectric layer fabrication followed standard photolithography procedures reported previously.^[45]^ Briefly, chromium-coated substrates with S1811 positive were exposed (11 s at 38-50 mW cm^-2^), developed in MF-321 developer, etched with CR-4 chromium etchant, and stripped with AZ-300T photoresist stripper. For the dielectric layer, the devices are placed under plasma oxygen (Harrick Plasma PDC-001, Ithaca, NY) for 1 min 30 s, after which they are immediately spin coated with a SU8-5 layer (10 s, 500 rpm, 30 s 2000 rpm), soft baked, and exposed to a sawtooth patterned mask to improve adhesion. After post-bake, substrates were developed in SU-8 5 developer. A final hard bake cycle was performed by ramping up to 180 °C in 15min, baking at 180 °C for 10 min and gradual cooling to room temperature. For the channel layer, soft-lithography procedure was followed. Si-wafers were cleaned with acetone/IPA and baked (150 °C, 15 min). The wafers were placed under plasma oxygen for 1 min 30s, after which they are immediately spin coated with SU8-2035 (500 rpm 10 s and 4000 rpm 30 s). The substrate was soft baked (55 °C 1 min, 75 °C 1 min, 95 °C 5 min) and exposed under a Quintel Q-4000 mask aligner (Neutronix Quintel, Morgan Hill, CA) (10 sec at 38-50 mW cm^-2^). Substrates were post baked (55 °C 1 min, 75 °C 1 min, 95 °C 5 min), and developed in SU-8 5 developer for 3 min 30 s upside down, without shaking. We followed a final hard bake cycle ramping up slowly to 165 °C for 10 min and cooling slowly to room temperature. The master mold was treated by chlorotrimethylsilane by vapour deposition in a desiccator for 45 min. PDMS (1:10 w/w ratio curing agent to prepolymer), poured over the mold and left to cure in an oven at 65 °C for 3 hours. Inlets and outlets were made using 0.75 mm or 0.35 mm biopsy punchers (World Precision Instruments, FL, USA), fitting 1/32″ OD tubing or 360 µm OD tubing respectively, after which the PDMS is carefully washed with IPA, DI water, and cleaned with tape to remove debris before device assembly. The PDMS slabs are treated with oxygen plasma for 30 s and directly aligned with the dielectric coated electrodes under a dissecting fluorescence microscope (Olympus IX73, 10X). Device channels were immediately treated with Novec 1720 fluorosilane polymer surfactant for 20 min, under a weight of 750 g. Devices were then air dried (20 min) and baked (150 °C, 30 min). To connect the droplet generator and the serpentine trap part of the device, a 2 cm piece of PEEK tubing (360 µm OD) was cut and treated with similar Novec 1720 treatment. Outlet blockers were made by hot gluing one end of a 1” PEEK 1/32″ OD tubing.

### Device operation

Gastight 500 μL glass syringes were prepared with the fittings and tubing as reported previously, adding one 2.5 mL syringe with a 1.87 mm magnetic stirring disk (V&P Scientific, San Diego, CA, USA). Syringes were installed on a low-pressure neMESYS pump system (Cetoni, Korbussen, DE), installed with a clamp holding a syringe stirrer (Nannostirus, V&P scientific, San Diego, CA, USA). The microfluidics device was placed inside a 3D printed pogo pin holder of which the base plate fits on the stage of an inverted microscope. The flow inside the microfluidic channel was observed under a 4X or 10X objective under bright-field illumination. Fluid flow and electrode actuation were controlled using an in-house automation system and user interface. In experiments that consisted of trapping, encapsulation, keeping and releasing, we followed a 5-step procedure (Supplementary Note 3, Supplementary Figure S4). First, priming was performed at a flow rate of 0.8 to 8 nL s^-1^ with PBS (Ca^2+^/ Mg^2+^ free) containing 2% Pluronics F-127 for 5 min, from inlet 1, to remove air bubbles and prevent cells sticking to the channel walls. The priming solution was then moved to outlet 1. The droplet generator was flushed with HFE-7500 with 2% fluorosurfactant, from inlet 4. Second, cells were loaded from inlet 1 at a flow rate of 1 nL s^-1^, in PBS (Ca^2+^/ Mg^2+^ free) or in complete media. Once cells are entering the inlet, the priming solution is turned off. We used MCF-7 cells as a model cell line and all reagents prior to operation were filtered sterilized. For trapping the cells, the filtered MCF-7 cells were resuspended in PBS (Ca^2+^ Mg^2+^ free) and pipetted into a UV sterilized 2.5 mL glass gastight syringe with stirring disk in a laminar hood. Third, droplets were generated using HFE 7500 with 2% w/v fluorosurfactant from inlet 4 with varying flow rates and PBS or media at 0.6 nL s^-1^ from inlet 3. On-demand droplet generation was performed by actuating electrodes with 15kHz 126 V_RMS_. Fourth, single-cell encapsulation can be performed. Droplet generation was paused after it stabilized, the cell flow was stopped, and the tubing was connected from outlet 3 to inlet 2, to the single-cell analysis chip. Phase change for encapsulation was performed using HFE 7500 with 2% w/v fluorosurfactant at a flow rate of 4 nL s^-1^. Fifth, under forward flow several operations (droplet releasing, keeping or merging) can be performed using electrode actuation patterns. To reverse the flow, the tubing in inlet 1 and outlet 1 need to be carefully removed, and a second oil syringe can be connected to outlet 2.

### Cell culture

MCF-7 cells were grown and maintained in DMEM containing 10% fetal bovine serum (FBS) with no antibiotics in an incubator at 37 °C with 5% CO_2_. Human lung squamous cell carcinoma dual-labeled (eGFP, Luciferase) stable NCI-H1299 (SCL-C01-HLG; Genecopoeia, Inc, Rockville, MD) cells were grown in RPMI-1640 containing 10% fetal bovine serum without antibiotics at 37°C with 5% CO_2_. For assays on device, cells were washed with PBS, trypsinized and resuspended in complete growth media. The cells were then centrifuged at 200 rcf for 4 min and the cell pellet was resuspended in either PBS or complete media without FBS to obtain an initial cell concentration of approximately 2 x 10^6^ cells mL^-1^. Cells were filtered through a 40 µm cell strainer (VWR) to remove cell clumps. An aliquot of the cell suspension (∼250 μL) was pipetted into a syringe for operation. Conditioned media for cell expansion was made by collecting complete growth media from 80% confluent NCI-H1299 cells (RPMI-1640 10% FBS, 1% penicillin/streptomycin) and adding 50% fresh complete growth media filter sterilizing and storing it at -80 °C until used.

### Viability assays

For viability assays, a 1X FDA/PI solution was made with 10 µL PI stock and 2.5 µL FDA stock, kept on ice, and used within 2 h. FDA/PI solution was placed in a 500 μL syringe covered with aluminium foil. After trapping of MCF-7 cells, the top two electrodes under each trap were actuated for 10 s (15 kHz 126 V_RMS_), after which the ground electrode was actuated for 10 s (15 kHz 126 V_RMS_). 1X FDA/PI was then flushed through the device at 1 nL s^-1^ and the device was incubated in the dark for 10 min. A similar viability test was also performed without electrode actuations, and in a 6-well plate for the same originating cell population used on the chip. Media was aspirated, and cells were washed with PBS. For the latter, 500 μL 1X FDA/PI solution was added to the well and cells were incubated in room temperature in the dark. After incubation, MCF-7 cells were imaged (FDA: *λ*_ex_ = 495 nm, *λ*_em_ = 520 nm, 300 ms exposure; PI: *λ*_ex_ = 535 nm, *λ*_em_ = 617 nm, 3 s exposure) under a fluorescent microscope and images were analyzed using ImageJ.

### H1299 Transfection and flow cytometry

pCRISPR_eGFP_314, pCRISPR_RAF1_94 and pCRISPR/Cas9_All_in_one_LacZ (Supplementary Information Table S3 and S4) were transformed into DH5*α* stocks, were grown overnight in LB with ampicillin, and purified (endotoxin free).

For lipid transfection of NCI-H1299 cells with an all-in-one pCRISPR/sgRNA plasmid in 24-well plates, on day 0, cells were subcultured in a 24-well plate to reach confluency the day after. On Day 1, cells were transfected using 5 µg DNA per well. After 48 h (Day 3), cells were harvested or subcultured into a 6-well plate. Confluent cells were trypsinized, centrifuged (200 rcf, 4 min), strained through a 40 µm filter, and resuspended in either PBS or flow cytometry buffer.

To obtain transfection efficiency, transfected and control cells were resuspended in sorting buffer (1X PBS (Ca^2+^/ Mg^2+^ free), 1mM EDTA, 25 mM HEPES pH 7.0, 1% FBS, sterilized using a 0.2 μm filter), were placed on ice, and loaded in a BD FACS Melody flow cytometer. Gating was performed for FSC and SSC on control population after which the transfected population was measured (561 nm laser).

### On-chip sorting, clonal recovery and expansion

For microfluidic sorting and cell recovery, PBS resuspended cells were loaded into a sterile 2.5 mL syringe with stir disk and 1/32″OD PEEK tubing. The device was primed with PBS 2% F-127, transfected NCI-H1299 cells were trapped at 4 nL s^-1^. One end of a sterile capillary tube was quickly dipped into HFE-7500 2% fluorosurfactant to render it hydrophobic. The capillary was placed on the outlet of the single-cell trapping device, a single droplets containing an isoclone was released individually. pCRISPR_eGFP_sg314 and pCRISPR_RAF1_sg94 transfected cells were sorted by forward releasing mCherry+/eGFP+ cell containing droplets on-demand. The oil flow to hydrodynamically release the remaining droplets to waste. In a biosafety cabinet, a droplet of 37°C 20µL conditioned media was placed on a hydrophobic PTFE membrane placed in a custom 3D printed holder. A single capillary containing the recovered droplet was immediately flushed with a total of 10 µL FC-40 oil, dispensing on top of the conditioned media droplet. After 1 min incubation, the droplet was recovered in a 96-well plate containing 50 µL warm conditioned media per well. After day 1, based on eGFP and or mCherry expression in wildtype, eGFP or RAF1 knockouts, single clones could be observed. After 5 days the media was changed to complete growth media. Clones were left to expand to 50% confluency in a 96-well plate at 5% CO_2_, 37°C in a humidified incubator. Clones were trypsinized and subcultured in 24-well plates and subsequently a 6-well plate, after which they were harvested for functional analysis or banking.

### Data analysis

Data analysis was performed with R v3.6.2. Data from Figure 3 was fitted with a 3 parameter logistic regression model with continuous predictors (Hosmer-Lemeshow goodness of fit test *p* > 0.05 for all three models) (Supporting Information Table S5). Image analysis was performed using Fiji by ImageJ. Fluid and electric field simulations were performed with COMSOL 5.4 Multiphysics (Supporting Information Note 2). Code was written in Python v2.7.15.

## 5. Acknowledgements

K.S. built the device, carried out experiments, analyzed the data and wrote custom software. K.S. and S.C.C.S designed the experiments. F.A. helped in device design, fabrication and characterization. A.B.V.Q. assisted in cell culture and transfection. G.S. aided in the design of the automation system software and hardware. K. S. and S. C.C. S. wrote the paper, and all authors reviewed the final version of the manuscript before submission. MCF-7 were generously donated by Dr. Alisa Piekny’s lab.

