## Supplemental Information for "One cell, one drop, one click: hybrid microfluidic mammalian single-cell isolation"

#### Supplementary Note 1: Device channel and electrode design

Optimizing single-cell trapping and encapsulation was done by designing the channel microfluidics device following hydrodynamic resistance ($R_h$) [1] and streamline based design rules[2].

$\Delta P = R_h Q$ , with $R_h$ the hydrodynamic resistance, $Q$ the volumetric flow rate and $\Delta P$ the pressure drop in the channel. The hydrodynamic resistance can be described as follows:

$$R_h = \frac{12\mu L}{wh^3(1 - 0.63\frac{h}{w})}$$

We consider three important hydraulic resistances: The trap constriction $R_2$, the trap $R_3$, and the bypass channel $R_1$ (Supplementary Figure S1).

$R_{eq} = \frac{R_1}{R_2+R_3}$ ,

Determining the width of the opening of the trap, the depth of the trap, the height of the channel, and the length of the bypass, we can calculate the three resistances. Before the cells are loaded, we aim for $R_2 + R_3 > R_1$ to encourage cell trapping. When a cell is trapped, we want $R_2 + R_3 \leq R_1$, $R_{eq} \approx 1$ to encourage $Q_1 > Q_{2+3}$ (or $Q_{bypass} > Q_{trap}$ ) while noting $R_3$ increases when a cell is trapped. Taking this into account, we next looked at the velocity streamline profile and positioned the trap near the curvature of the main channel (i.e. the end of a serpentine channel) along with narrowing the width of channel near trap to improve cell trapping.

For the electrode design, four electrodes were sufficient to perform desired operations. A common ground electrode wire going through the center of the trap was chosen to act as a ground in case all four electrodes require a potential. Interdigitation was used and electrodes have a minimum gap of 14 µm. The wiring has a thickness of 20 µm. The amount of wiring going under or through channels is minimized, as actuations through wiring can manipulate the flow. This limited us to only equip 6 out of 12 traps with 4 electrodes each.

### Supplementary Note 2: Computational fluid dynamics and electric field simulations

The model portraying the channel geometry with flow velocity profile, velocity streamlines and pressure was stationary simulated using COMSOL Multiphysics 2D creeping flow physics. Inertia and turbulence were neglected, and no slip was set for channel walls. The fluid within the channels was PBS, the boundaries were PDMS.

Conditions used for modeling:

P = 0 Pa for outlet pressure

$\boldsymbol{u} = \begin{cases} x = 0 \\ y = 0 \end{cases} \; m\, s^{-1}$ for initial velocity profile

$V_0 = 0.0008$ µL $s^{-1}$ for volumetric laminar inflow

The model portraying the electric field generated by different actuation patterns, was simulated in COMSOL Multiphysics with a static electricity module (Supplementary Figure 6). Electrodes were modeled as 2D surfaces without thickness, under a 7 µm SU-8 surface, covered with an HFE oil of 30 µm thickness (Figure 1). We did not place PDMS features on top.

Conditions used for modeling the electric field above the dielectric:

T= 293.15 K is the temperature

As potential $V_0$:

1.8 $V_{pp}$ = 142$V_{RMS}$ for droplet generation

1.6 $V_{pp}$ = 126 $V_{RMS}$ for droplet release, keep and encapsulate

And with ground potential and initial potential V= 0 V

The model was static, stationary and materials were continuous. For each model, the not actuated electrodes were selected as ground.

Under static conditions, the electric potential, $V$, is defined by the relationship

$$E = -\nabla V$$

and the electric displacement or flux density $\boldsymbol{D}$ is defined by the relationship

$$\mathbf{D} = \varepsilon_0 \mathbf{E} + \mathrm{P}$$

with the electric constant $\varepsilon_0$ , electric field $\boldsymbol{E}$, and P.

Following Gauss' Law:

$$-\nabla(\varepsilon_0 \nabla \mathrm{V} - \mathrm{P}) = \rho$$

With electric field $\boldsymbol{E}$, tangential to the xy-plane (3D) this becomes:

$$-\nabla\, \mathrm{d}\, (\nabla \mathrm{V} - \mathrm{P}) = \rho$$

with d and ρ the total electric charge density.

In COMSOL, the magnitude of the electric vector field (V $m^{-1}$) is calculated with:

$$\boldsymbol{E} = \sqrt{{E_x}^2 + {E_y}^2 + {E_z}^2}$$

**Table S1. Additional device statistics**

| Dispersed | Continuous | Contact angle (°) | Standard deviation | N |
|---|---|---|---|---|
| PBS | HFE 7500 + 2% Ran | 139.55 | 4.9 | 10 |
| DI Water | Air | 128.968 | 4.53 | 10 |
| **Droplet volume** | | **Volume (pL)** | **Standard deviation** | **N** |
| After single-cell encapsulation | | 150.25 | 55.58 | 11 |
| On-demand droplet-generation | | 250.9 | 39 | 44 |

**Table S2. Parameters for multi-physics modeling**

| Property | Value |
|---|---|
| SU8-5 Relative Permittivity | 4.5 |
| PDMS Relative permittivity | 2.75 |
| PDMS density | 970 [kg $m^{-3}$] |
| PBS density | 1000 [kg $m^{-3}$] |
| PBS dynamic viscosity | 0.0008882 [Pa s] |
| HFE 7500 oil relative permittivity | 5.8 |
| HFE 7500 oil boundary size | 500x 500 [$\mu m^2$ ] |
| Dielectric thickness SU-8 5 | 7.0 [µm] |
| SU-8 5 boundary size | 500 x 500 [$\mu m^2$ ] |
| Electrode sizes | 110 x 110 and 110 x 40 [$\mu m^2$ ] |
| Ground electrode size | 10 [µm] |
| Electrode gap width | 14 [µm] |

### Supplementary Note 3: Automation setup and device operation

The microfluidics device is placed inside a 3D printed pogo pin PCB board holder of which the base plate fits on the stage of an inverted microscope (Olympus IX7) [3]. The flow is observed under a 4X or 10X objective under bright-field illumination. A neMESYS Low Pressure pump with five syringe units, and an Arduino Uno were connected to a PC, and operated through Python 2.7. The Arduino Uno is connected to a stack of 120 optocouplers - powered by a 5 $V_{DC}$ power supply. A 15 kHz sinusoidal signal (function generator) is amplified. The Python protocol is executed and the GUI is started after which flows are started with disconnected tubing to remove air bubbles in the system. Tubing is carefully connected to the device (Supporting Information Figure 3).

Priming is done from inlet 1 with PBS containing 2% Pluronics F-127, filled in a 500 µL gaslight syringe. After the device is primed (for at least 5 min), the priming solution is moved from the inlet to either outlet and turned to a low flowrate. Cells are loaded from inlet 1, either re-suspended in PBS or their original media, and placed in a 2.5 mL syringe with a 7 mm magnetic stirring disk. The syringe is stirred continuously throughout the procedure. Once cells are entering the device, and leaving through inlet 2, the flow of the priming solution can be stopped and cells will enter the trapping array. Phase change for single-cell encapsulation is performed using HFE 7500 with 2% fluorosurfactant arriving from the droplet generator with Inlet 4 connected to a 500 µL syringe. Inlet 3 is connected to a 500 µL syringe with an aqueous solution (droplet content). Outlet 3 is connected to Inlet 2 with PEEK tubing. Electrodes are actuated using the GUI. Once the cells are encapsulated, additional droplets can be generated on-demand using the GUI, by using HFE 7500 with 2% fluorosurfactant at Inlet 4 with varying flow rates, and aqueous flow from Inlet 3 at 0.6 nL $s^{-1}$. In- or out-lets can be blocked using PEEK tubing with glued ends, if needed.

**Supplementary Note 4: Device assembly and channel treatment**

Novec 1720 contains a fluorosilane polymer surfactant dissolved in an ether solution. The solution evaporates and is an ideal solution to use to avoid clogging in the trap or channel (which is sometimes not the case when using oil-based surfactants). Clean PDMS channel slabs are treated with oxygen plasma for 30 s and directly aligned on top of a clean dielectric coated electrode patterned glass, under a microscope (4X). The device is sealed with transparent adhesive tape and pressure is applied to ensure strong adhesion. Immediately, the device channel is treated with Novec 1720 for 20 min. Device was air dried for 20 min. Bake device at 150 ºC for 30 min, while applying 750 g weight on top of the device. For the droplet bridge, a 2 cm piece of PEEK tubing is treated with Novec 1720 for 20 min. To reuse devices, flush the device with Fluorinert FC-40 to remove the oil containing surfactant in the traps, and then bake at 100 ˚C for 2 h. If actuations were applied, the device was washed with FC-40, and treated with Novec 1720 (similar to the protocol before). Chips that were not used for 14 days should be treated again with Novec 1720. If needed, chips can be washed with a 1% Triton X-100 solution to flush out debris, and then continue with an FC40 wash.

**Table S3. Cells and plasmids**

| Cells Lines | Transgene integration | Source |
|---|---|---|
| NCI-H1299 | Luciferase, eGFP, KanR | Genecopeia SL001 |
| MCF-7 | N/A | Donated by Dr. Piekny's Lab |
| **Plasmids** | **Relevant characteristics** | **Source** |
| pCRISPR_eGFP_497 | AmpR, Neo/KanR | Addgene #111824 |
| pCRISPR_RAF1_94 | AmpR, Neo/KanR | Addgene #111824 |
| All_in_one_CRISPR/Cas9_LacZ | AmpR | Addgene #74293 |

**Table S4. Primers**

| Name | Purpose | Orientation | Sequence |
|---|---|---|---|
| eGFP_314_2919 2938 | sgRNA | Forward | CTTCTTCAAGTCCGCCATGC |
| eGFP_314_3307 3326 | sgRNA | Reverse | ATGTGATCGCGCTTCTCGTT |
| RAF1_16_11 31 | sgRNA | Forward | CAGTAGGGCTCGCGCAGAATC |
| RAF1_16_465 484 | sgRNA | Reverse | CCCTGGAAGCCGATCCCTCA |

**Table S5. Three parameter logistic regression model.** Parameters and the Hosmer-Lemeshow goodness of fit statistic of each fitted model.

$$y = \frac{A}{1 + e^{\frac{B-x}{C}}}$$

| Data | Asymptote (A) | Inflection point (B) | Scale (C ) | P (GOF) |
|---|---|---|---|---|
| Forward flow on-demand droplet release | 1.1646 | 3.23 | -1.92 | 0.9888 |
| Reverse flow Hydrodynamic droplet release | 0.96059 | 5.68 | 1.73 | 0.9616 |
| Reverse flow on demand droplet keeping | 0.80613 | 64.01 | -5.87 | 0.9967 |

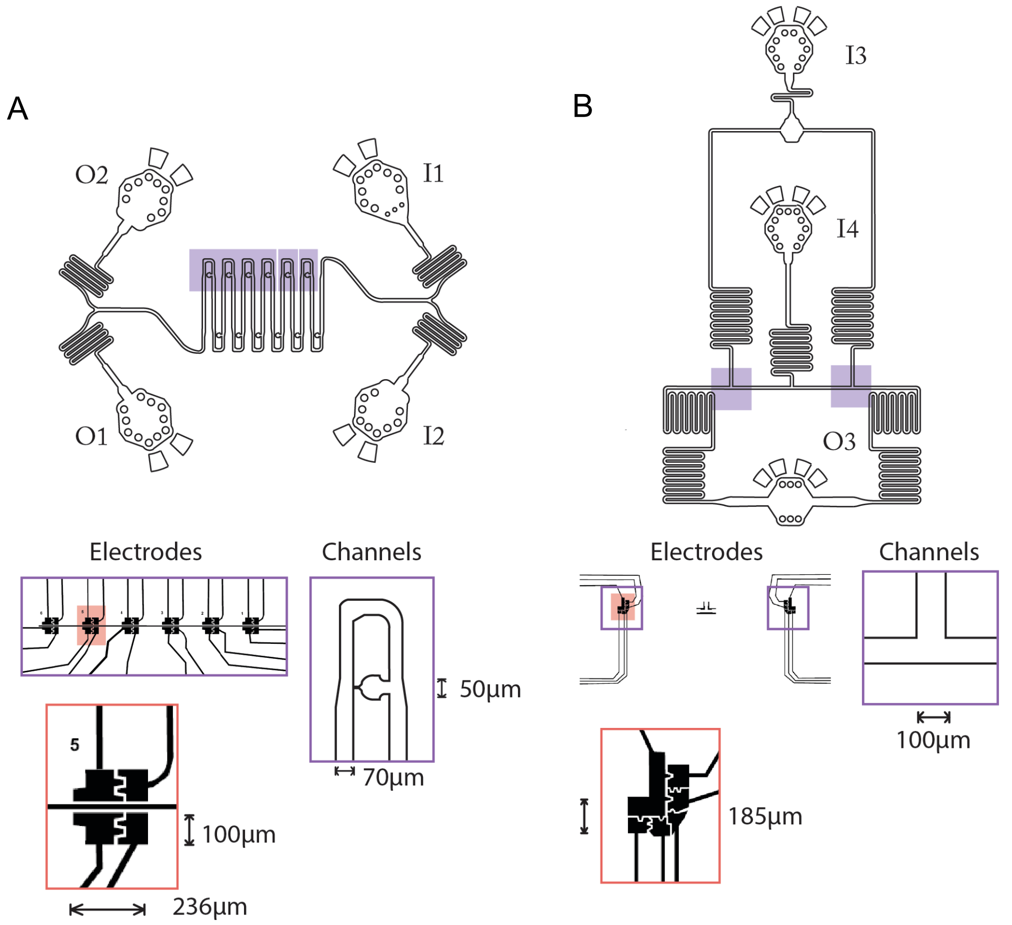


**Supplementary Figure 1 - Device electrode and channel geometry.** The device consists of of two separate PDMS layers, a serpentine trapping channel and a droplet generator. A) The serpentine trapping channel contains two inlets (I1, I2) and two outlets (O1, O2). The channel width is 70µm with a bypass channel width of 50 µm, and height of 35 µm. Highlighted is the six traps and is aligned with electrodes below the trap (enlarged). B) The droplet generator has two inlets (I3, I4) and one outlet (O3). The channel width is 100 µm, and height is 35 µm. Highlighted area is lined with electrodes (enlarged).

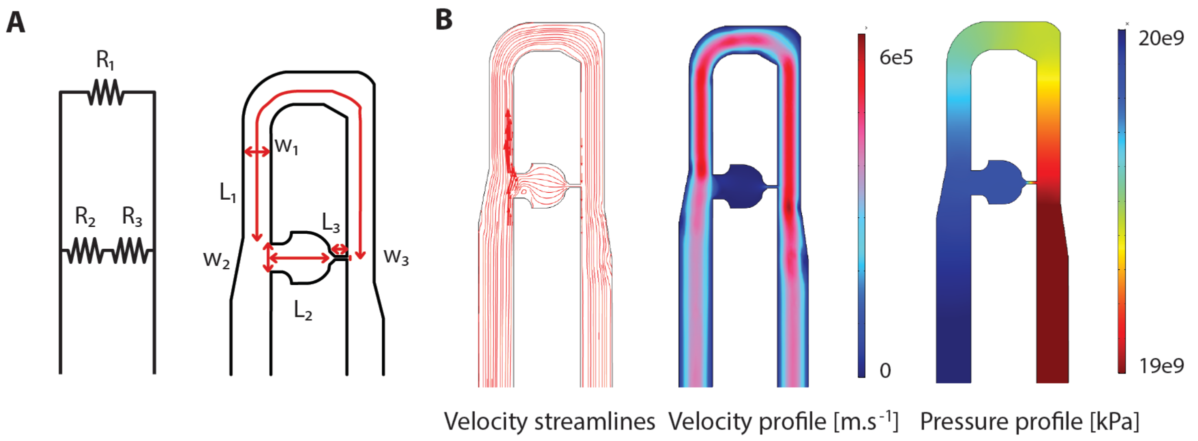


**Supplementary Figure 2 Resistance based channel design**. A) An equivalent resistor circuit for the bypass and trapping area on the device. Hydrodynamic resistance in the cell trap ($R_3$) will increase upon trapping a cell. The flow prefers the path of least resistance and the bypass path ($R_1$) is preferred. The ratio of hydrodynamic resistance in bypass on trap hydrodynamic resistance is used for designing an efficient cell trapping and phase change system. The length and width of the bypass ($L_1$ =1000 µm $w_3$ = 50 µm) and trap (L = w = 50 µm) were modified to obtain higher efficiencies. Dimensions are as indicated. B) Numerical simulations with COMSOL Multiphysics show flow velocity pattern across trapping array.

A

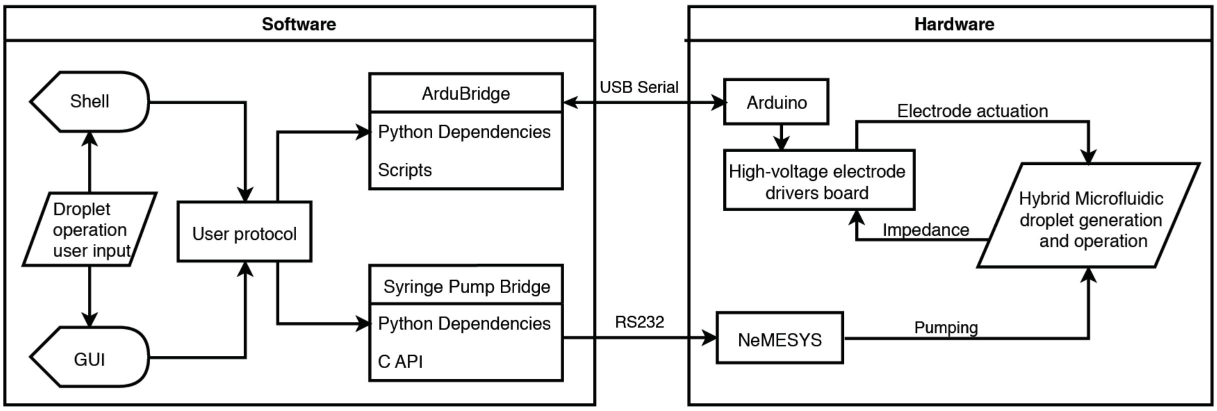


B

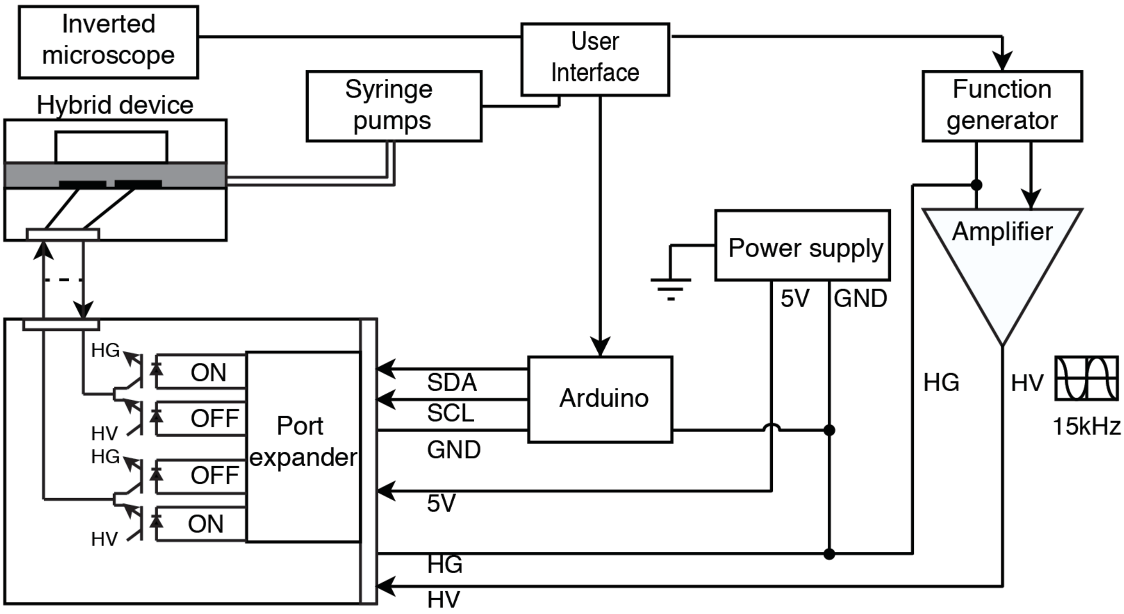


**Supplementary Figure 3 - Software and hardware diagram.** A) Unified Modeling Language style diagram showing communication lines between software and hardware, and with integration of the pumps and electric potential operation through Python 2.7. Through a GUI, the user can perform on-demand droplet operations such as droplet generation, encapsulation, keeping or releasing a droplet. For example, the user can create any number of droplets with a set time interval or can perform on-demand encapsulation in any of the traps or release or keep the droplet simply by the click of a button. The GUI accesses a Bridge that writes to an Arduino or a syringe pump system. B) Hardware setup. Arduino controls an $I^2C$ communication protocol to address specific optocouplers. Automation system hardware setup is similar to previously reported.[3]

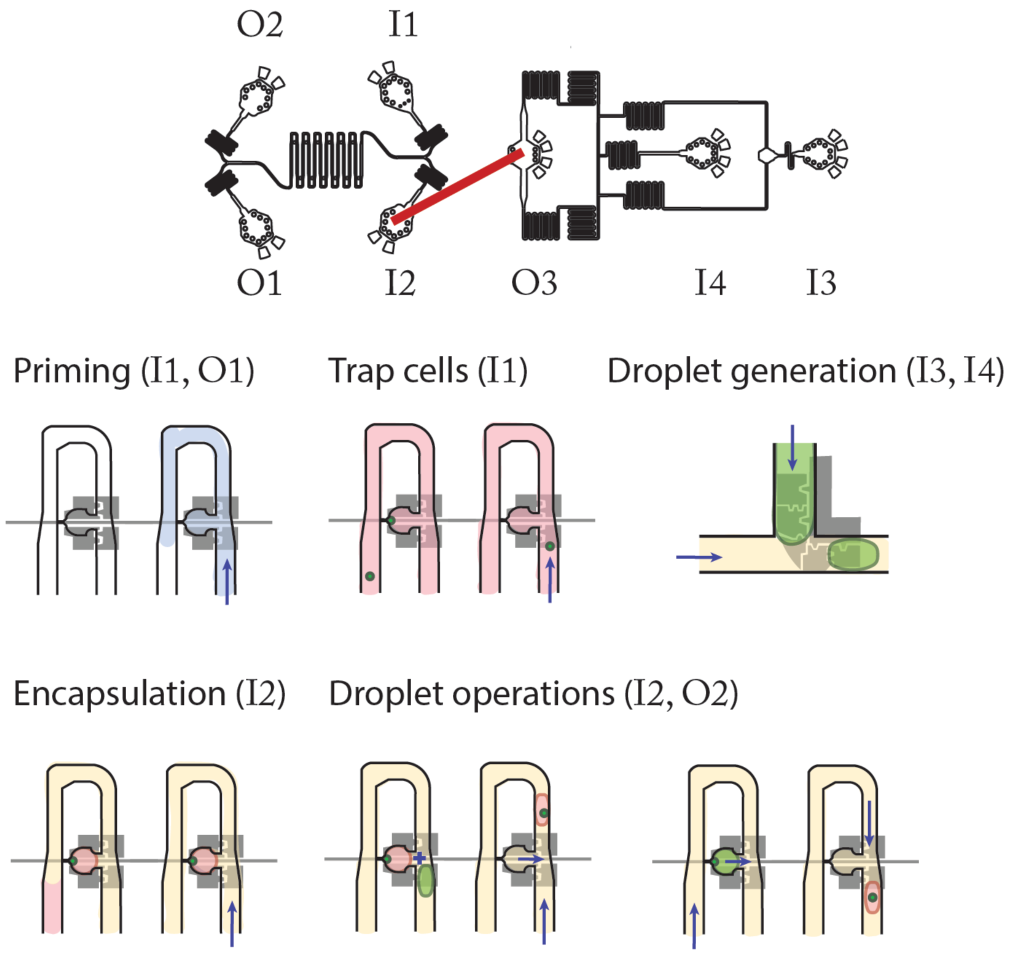


**Supplementary Figure 4 - Workflow of device operation.** Priming the device with PBS and 2% Pluronics F-127 for 5 min. MCF-7 cells in PBS are trapped. Droplet generation is started and stabilized, after which the aqueous flow is stopped. When all traps are loaded, oil (HFE 7500 2% Ran fluorosurfactant) is loaded at 4 nL $s^{-1}$ by connecting the droplet bridge. Oil flow shears off a small volume of remaining PBS, which forms a droplet around the cells. Droplets are brought in through the droplet bridge and droplet operations can be performed. Oil flow can be reversed to collect droplets.

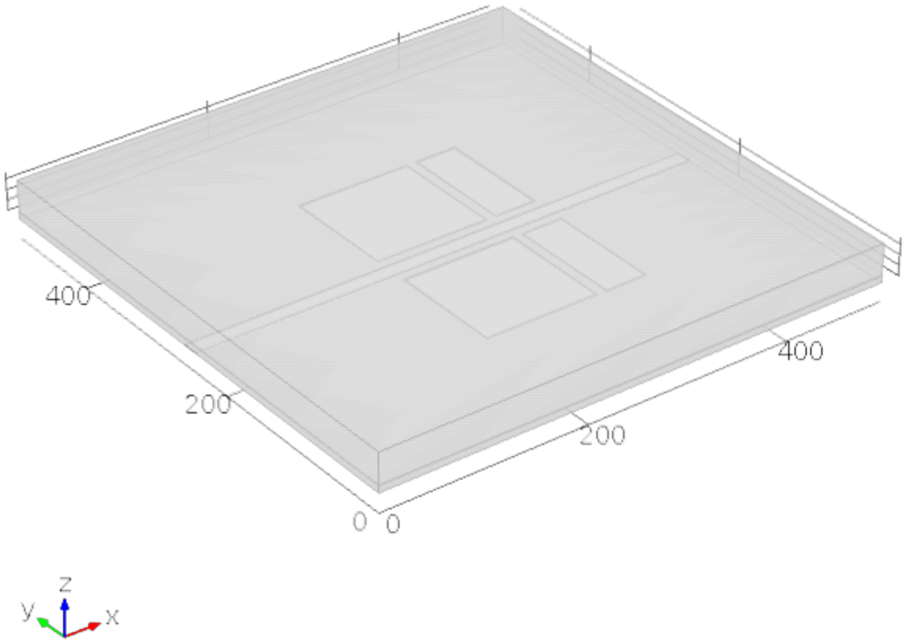


**Supplementary Figure 5 - Geometry used for electrostatic field calculations.** Geometry used for electric potential (V) and electric field (V $m^{-1}$) modeling on a 7 µm SU8-5 dielectric layer above co-planar electrode surfaces using COMSOL Multiphysics electrostatics numerical modelling. Dimensions in the image are in µm. Top layer is a 35 µm HFE 7500 oil layer, under which a 7 µm thick SU-8 5 layer is positioned with defined areas of potential or grounding.

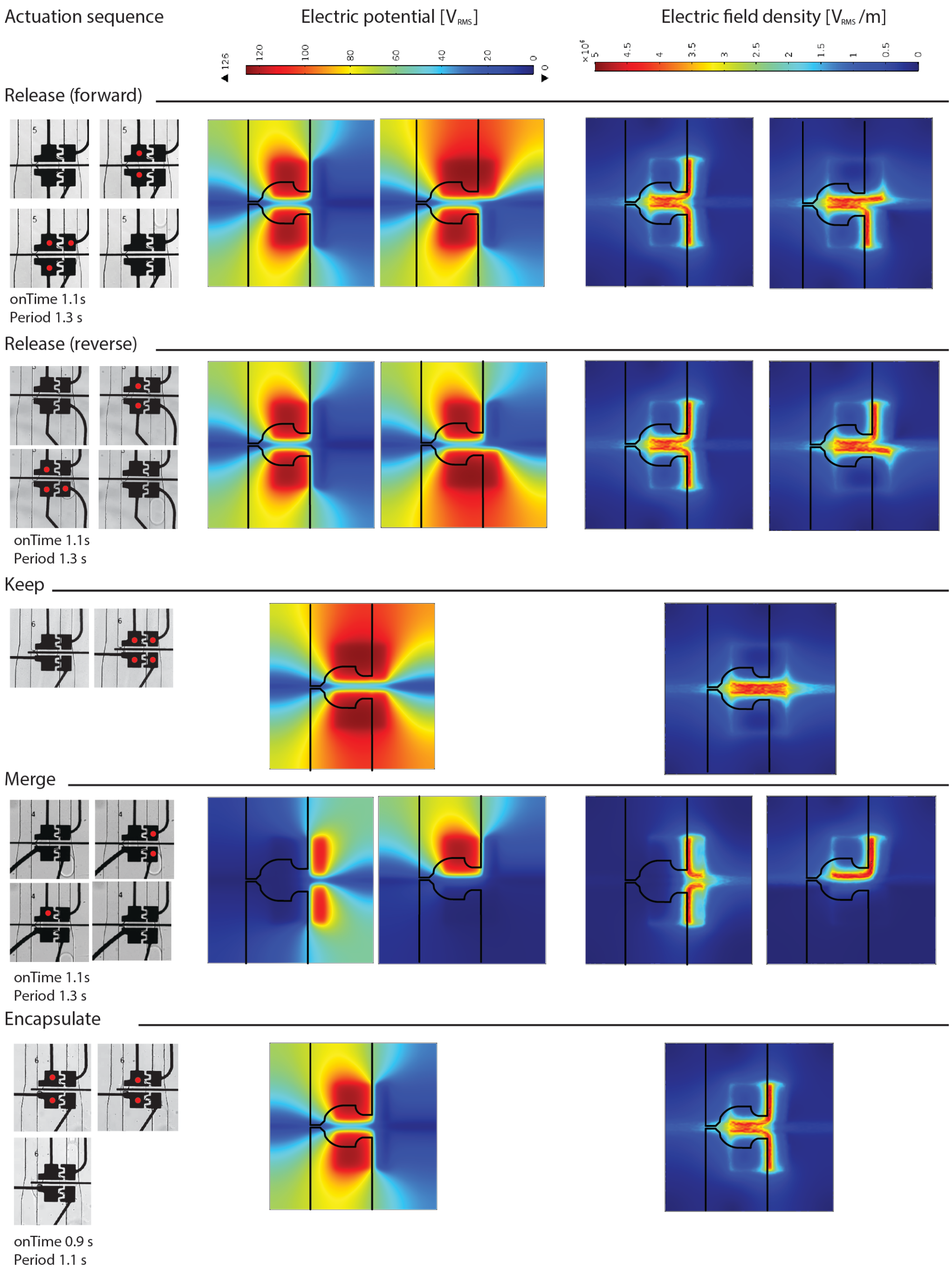


**Supplementary Figure 6 - Electrode sequences of on-demand droplet operations**. Electric potential (V) and electric field (V m$^{-1}$) on 7µm SU8-5 dielectric layer above co-planar electrode surfaces were modelled using COMSOL Multiphysics electrostatics numerical modelling. Actuated electrodes are marked by a red dot (bright field image, 15X). Droplet release takes place under 15 kHz 126 $V_{RMS}$ with a pulse width of 1.1 s and period 1.3 s. Droplet keeping takes place under 15kHz 126 $V_{RMS}$ with varying pulse width. Droplet merging takes place under 15kHz 126 $V_{RMS}$ with pulse width of 1.1 s and period 1.3 s. Encapsulation takes place under 15 kHz 126 $V_{RMS}$ with pulse width of 0.9 s and period 1.1 s.

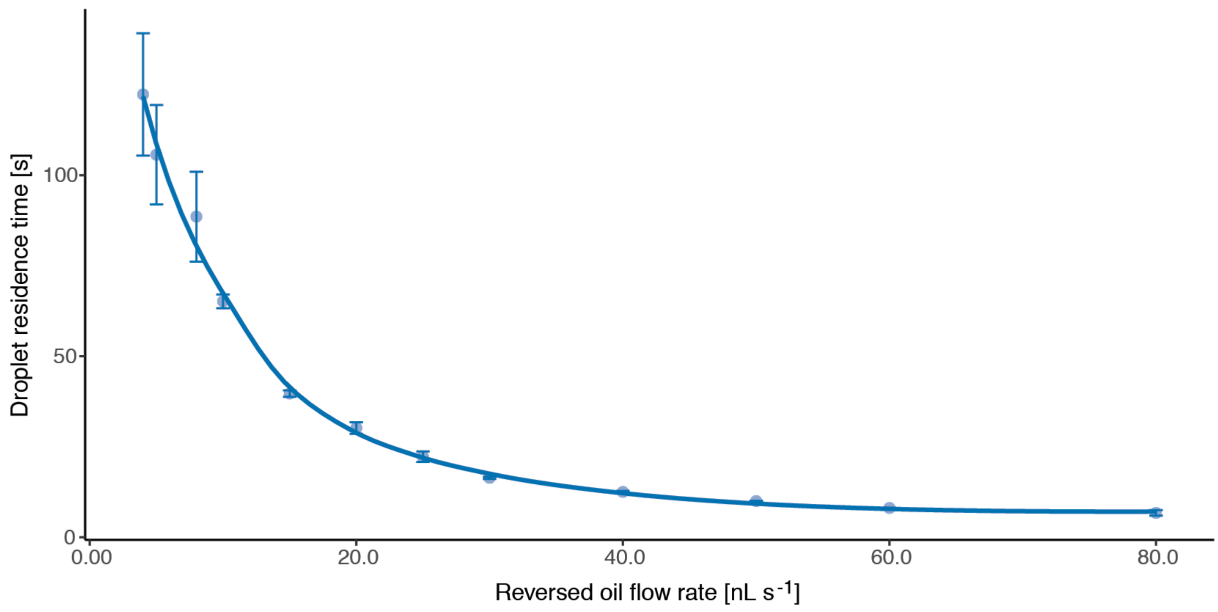


**Supplementary Figure 7 - Droplet residence time.** A graph showing the residence time for the last droplet (trap 6) to leave the trapping array under different flow rates. At low flow rates (< 10 nL $s^{-1}$), residence time is highly variable. At high flowrates (> 10 nL $s^{-1}$), the time required to keep droplets on chip is shorter (requiring less actuation time).

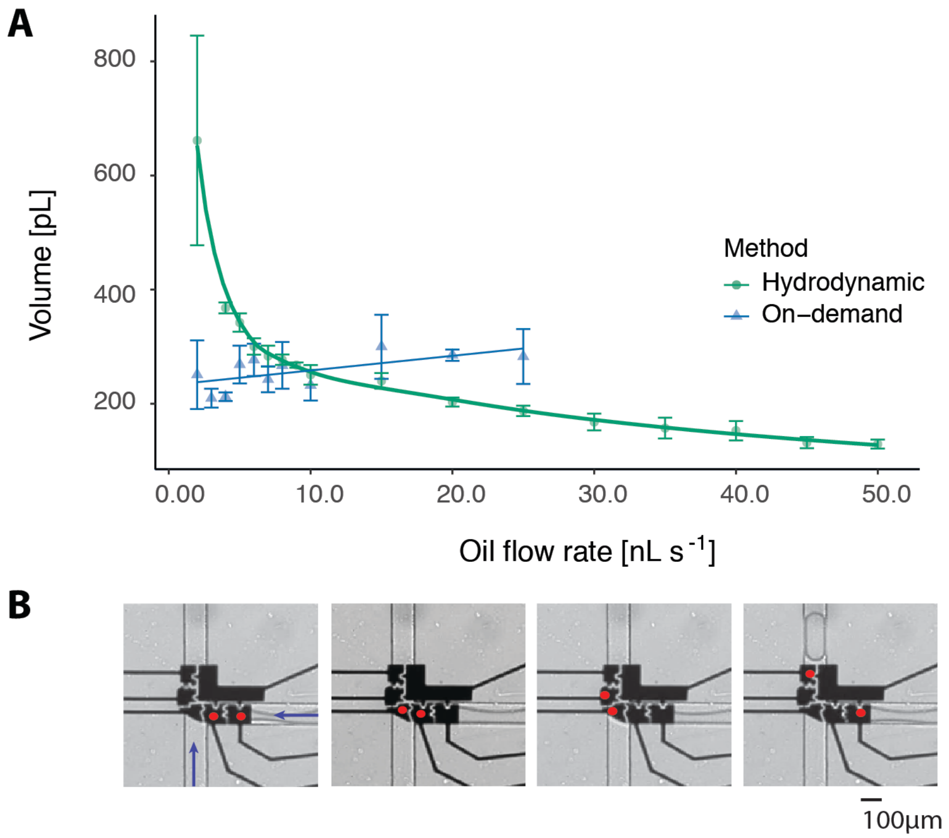


**Supplementary Figure 8 - Characterization of on-demand droplet generation.** A) Electrodes are actuated in a serial fashion to generate droplets on-demand under 15 kHz and 126 $V_{RMS}$ (bright field image, 10X). B) Average droplet volume (in pL) of on-demand generated and hydrodynamically generated droplets. Droplets are generated with a double T-junction of 100 µm width. On-demand droplet-generation shows stable droplet volume for on-demand droplet generation of 207.5 pL droplets. Droplet volume was estimated by multiplying droplet area and a channel height of 35 µm using Fiji (ImageJ).

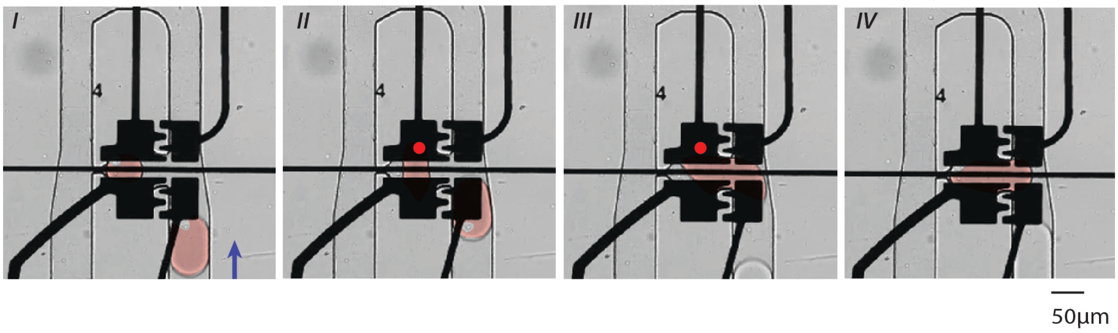


**Supplementary Figure 9 - Droplet merging**. Four images showing the process of droplet merging. In Frame 1, a droplet with a single MCF-7 cell is inside a trap and a droplet containing a cell is generated and in the main channel flowing at a rate of 4 nl $s^{-1}$. In Frame 2, an electrode (shown by the red dot) is activated to initiate the merging process. In Frame 3, the droplets are merged and can be stored in the trap. In Frame 4, the merged droplet can be kept or released via on-demand functions.

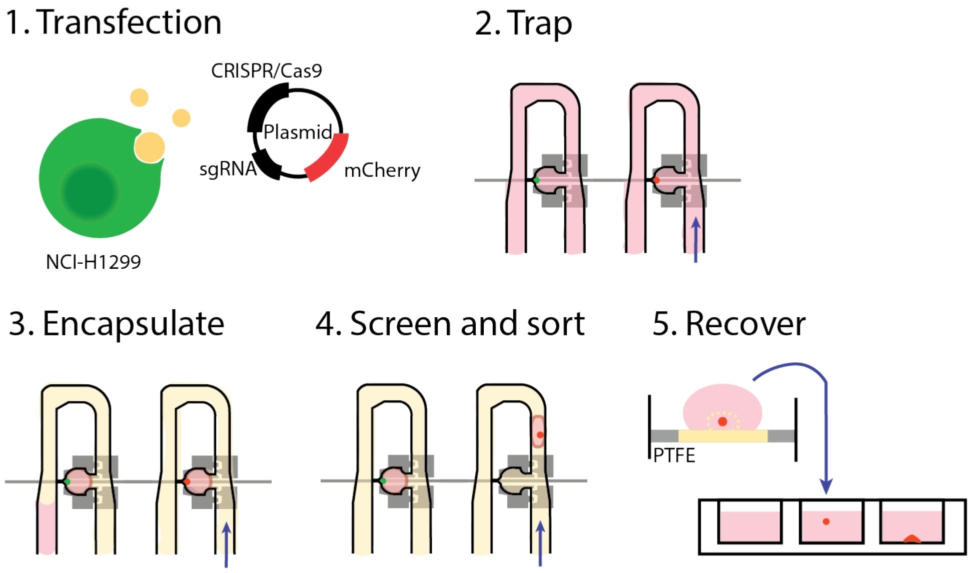


**Supplementary Figure 10 - Overview of hybrid microfluidics isoclonal recovery pipeline**. After transfection (1), the device is loaded with a heterozygous transfected cell suspension and single isoclones are trapped (2). An HFE-7500 2% Ran surfactant oil flow is flown through the device and electrodes are actuated in order to encapsulate single-cells in droplets (3). Single isoclone containing droplets can be selected and released on demand. After capillary recovery and centrifugation, isoclones are recovered in 96-well plates and maintained for expansion (4).

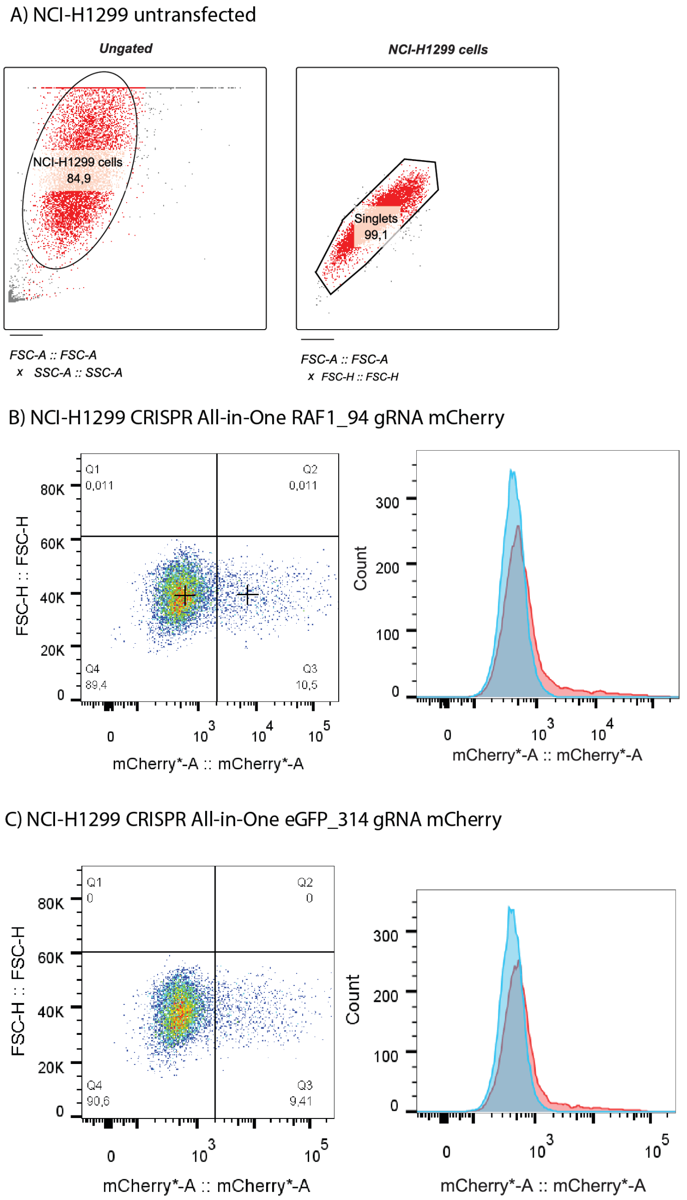


**Supplementary Figure 11 - Transfection efficiency determined by flow cytometry.** A) Gating of untransfected NCI-H1299 cell singlets based on cell morphology to remove dead cells. Flow cytometry of NCI-H1299 cells that have been transfected by B) a pCRISPR_RAF1_94 plasmid that contains a sgRNA targeting the RAF1 gene or C) a pCRISPR_eGFP_497 plasmid that contains a sgRNA targeting the eGFP gene. (Left) Forward scatter versus mCherry expression plots that are gated at a fluorescence intensity of $10^3$ after 48 h of transfection. A transfection efficiency of 10.5% and 9.41% is observed for the RAF1 and eGFP plasmid respectively. (Right) Histograms showing transfected single cells (red) and non-transfected single cells (blue).

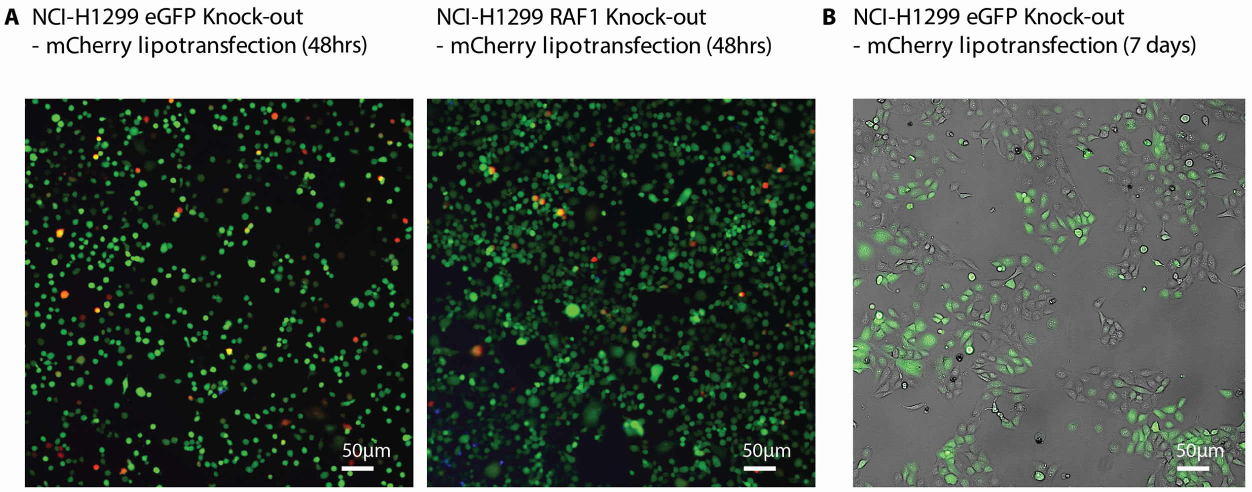


**Supplementary Figure 12 - Microscopy images of transfected NCI-H1299.** A) lipid mediated transfection with eGFP targeting and RAF1 targeting sgRNA encoding plasmid. Fluorescent microscopy image of NCI-H1299 lung squamous cell carcinoma's after 48hrs of lipid-mediated transfection, showing mCherry expression ($\lambda_{ex}$: 585 nm/$\lambda_{em}$: 608 nm), native eGFP expression ($\lambda_{ex}$: 488 nm /$\lambda_{em}$: 509 nm) and uptake of ViaQuant™ Blue Fixable Dead Cell Stain ($\lambda_{ex}$: 350 nm /$\lambda_{em}$: 440 nm). B) Phenotypic eGFP loss visible in heterozygous eGFP KO strain after 7 days in culture.
